# A novel variant of the *Listeria monocytogenes* type VII secretion system EssC component is associated with an Rhs toxin

**DOI:** 10.1101/2023.02.17.528482

**Authors:** Kieran Bowran, Stephen R. Garrett, Arnoud H. M. van Vliet, Tracy Palmer

## Abstract

The Type VIIb protein secretion system (T7SSb) is found in Bacillota (firmicute) bacteria and has been shown to mediate interbacterial competition. EssC is a membrane-bound ATPase that is a critical component of the T7SSb and plays a key role in substrate recognition. Prior analysis of available genome sequences of the foodborne bacterial pathogen *Listeria monocytogenes* has shown that although the T7SSb was encoded as part of the core genome, EssC could be found as one of seven different sequence variants. While each sequence variant was associated with a specific suite of candidate substrate proteins encoded immediately downstream of *essC*, many LXG-domain proteins were encoded across multiple *essC* sequence variants. Here we have extended this analysis using a diverse collection of 37,930 *L. monocytogenes* genomes. We have identified a rare eighth variant of EssC present in ten *L. monocytogenes* Lineage III genomes. These genomes also encode a large toxin of the rearrangement hotspot (Rhs) repeat family adjacent to *essC8*, along with a probable immunity protein and three small accessory proteins. We have further identified nine novel LXG-domain proteins, and four additional chromosomal hotspots across *L. monocytogenes* genomes where LXG proteins can be encoded. The eight *L. monocytogenes* EssC variants were also found in other *Listeria* species, with additional novel EssC types also identified. Across the genus, species frequently encoded multiple EssC types, indicating that T7SSb diversity is a primary feature of the genus *Listeria*.

**DATA SUMMARY:** All genome sequences used in this study are available via Genbank, and the assembly accession numbers are provided in Table S1. This file also lists relevant metadata (name, source category, country, year and clonal complex).

**IMPACT STATEMENT:** *Listeria monocytogenes* is a soil-borne saprophytic bacterium and a food-borne pathogen of humans. Decomposing plant matter and the human GI tract are rich in diverse microbial species and to colonise these niches *L. monocytogenes* must be able to compete with other bacteria. The type VII secretion system (T7SS) of Bacillota has been shown to secrete protein toxins that target other bacteria. In this study we have analysed a diverse collection of *L. monocytogenes* genome sequences to study the diversity of the *Listeria* T7SS and its putative effector proteins. We show that the EssC component of the *L. monocytogenes* T7SS is highly diverse, clustering into one of eight sequence variants. Each EssC variant is associated with a specific toxin candidate, and the EssC8 variant T7SS likely secretes a novel rearrangement hotspot (Rhs) repeat toxin. We also identify multiple new LXG-families of T7SS toxins and describe genomic hotspots where they are encoded. We find no link between EssC variants and clinical outcome. In agreement with this, analysis of EssC variability in available genomes of other *Listeria* species showed that all eight *L. monocytogenes* EssC variants are present in non-monocytogenes *Listeria* species.

## INTRODUCTION

In their natural environments, bacteria live in complex communities with other microbes, viruses and often, higher eukaryotes. In order to co-exist, bacteria have developed numerous strategies, including competitive and co-operative mechanisms e.g. (1, 2). A major way that bacteria interact with biotic and abiotic factors is through the secretion of extracellular proteins, and to date 11 protein secretion systems have been described (3). While most of these are unique to Gram-negative bacteria, the Type VII secretion system (T7SS) is found in many monoderm Gram-positive bacteria, including Staphylococci and Streptococci, and in diderm Gram-positives such as Mycobacteria (4, 5).

There are commonalities and differences between the T7SS of Mycobacteria and of Bacillota such as *Staphylococcus* spp., resulting in them being assigned T7SSa (actinobacteria) and T7SSb (Bacillota) (6, 7). One of the key similarities is the requirement of a membrane-bound AAA+ ATPase for secretion activity. In T7SSa, the ATPase is named EccC and is arranged as a hexamer at the centre of a 2.3 MDa membrane-bound protein complex (7–9). The corresponding ATPase in T7SSb is termed EssC, and shares sequence similarity with EccC but additionally has two forkhead associated domains at its N-terminus that are essential for activity (10, 11). A second common feature between the two systems is the presence of helical hairpin substrate proteins from the WXG100 family. These small proteins form homo-(T7SSb) or heterodimers (T7SSa) and are secreted in a folded state (12, 13). Other substrate families appear to be unique to either of the two systems, with the T7SSa secreting PE/PPE and Esp proteins, whereas T7SSb substrates have an LXG or YeeF-like domain (14–17).

Growing evidence indicates that Bacillota employ their T7SSb for competitive interactions with other bacteria. *Staphylococcus aureus* secretes a YeeF-like domain-containing DNase, EsaD, and an LXG-domain membrane depolarising toxin, TspA, which can target closely related bacteria (17, 18). Four LXG toxins have been identified in *Streptococcus intermedius*, and LXG toxins have also been shown to mediate bacterial antagonism in *Enterococcus faecalis* and *Bacillus subtilis* (16, 19–21). However, to date, the largest number of candidate T7SSb toxins have been identified in *Listeria monocytogenes* (22).

*L. monocytogenes* is a food-borne pathogen that is found in water, soil and decomposing vegetation (23). From here it can make its way into to food chain, where its ability to resist low pH and high salt contribute to its survival in food. Once ingested, it can cause numerous diseases from abortion in pregnant individuals to bacteraemia and death in up to 30% of cases (24, 25). *L. monocytogenes* genomes have been classified into one of four evolutionary lineages and multiple clonal complexes, with Lineages I and II most frequently associated with human disease (26). Genes encoding the T7SSb are found in all sequenced genomes to date, and are located in a region of high sequence variability (27).

Our prior genomic analysis revealed that the gene encoding the T7SSb core component, EssC, is found as seven different variants across *L. monocytogenes* genomes. The sequence variability corresponds to the final two ATPase domains of EssC, and each EssC variant is adjacent to a variable region (termed variable region 1) that likely encodes one or more secreted toxin substrates specific to each EssC variant (22). We found that the EssC1 variant is the most prevalent and that *essC1*-specific toxins have a YeeF-like domain, related to *S. aureus* EsaD, while *essC5*, *essC6* and *essC7* genomes encode LXG-domain toxins at the equivalent position (22). In addition to EssC-specific suites of substrate proteins, we also found other genomic hotspots where T7SSb toxins could be encoded, identifying 40 different LXG toxins in sequenced genomes. Collectively this high diversity implies that the T7SSb plays a key role in bacterial antagonism.

Here we further investigate *L. monocytogenes* T7SSb variability through analysis of a collection of 37,930 *L. monocytogenes* genomes obtained from public repositories. We have identified a rare eighth EssC variant unique to Lineage III strains, which is genetically associated with a likely antibacterial toxin of the rearrangement hotspot (Rhs) repeat protein family. We also describe further LXG candidate toxin proteins across the species and four additional chromosomal ‘hotspots’ where they may be encoded. This high variability in EssC type is observed in other *Listeria* species, suggestive of competitive strategy that is widely utilised across the genus.

## METHODS

### Generation of a *L. monocytogenes* genome repository

The NCBI Pathogen Detection database (https://ncbi.nlm.nih.gov/pathogens) was used to download assembly accession numbers and metadata of assembled *L. monocytogenes* genomes of Lineages I-IV. Genomes were downloaded with ncbi-genome-download version 0.3.1 (https://github.com/kblin/ncbi-genome-download), and assembly metrics were checked with Quast version 4.6 (28). The 7-gene multilocus sequence types (MLST) and clonal complexes (CC) for *L. monocytogenes* (29) were determined using mlst version 2.23 (https://github.com/tseemann/mlst), while core genome MLST (cgMLST) was performed using chewBBACA version 2.8.5 (30) and the 1,748 allelic scheme from the Listeria PubMLST site at Institute Pasteur (https://bigsdb.pasteur.fr/listeria) (31). Phylogenomic trees were built from cgMLST profiles using Grapetree version 1.5 using the RapidNJ algorithm (32). The list of *L. monocytogenes* genomes included, MLST sequence type, clonal complex, order in cgMLST trees, accession numbers and associated metadata are provided in Table S1. Isolates were assigned as clinical when obtained from human samples, while assigned non-clinical when obtained from food, factory or environmental sources. Those obtained from animal sources were not assigned clinical or non-clinical as this could not be determined from the available information.

### Querying the *L. monocytogenes* genome repository for EssC and LXG toxin variants

The *L. monocytogenes* genome repository was queried using Abricate version 1.0.1 (https://github.com/tseemann/abricate), using DNA sequences representing conserved and variable regions of the genes encoding the EssC1-8 proteins and the LXG proteins D, E, F, G and Omicron. All genomes were also converted into predicted open reading frames using Prodigal version 2.6.3 (33), and screened using amino acid translations of the conserved and variable regions using BLAST+ version 2.13.0 (34).

### *L. monocytogenes* genomes and annotation for analysis of the *essC8* locus

*L. monocytogenes* EGDe (NC_003210.1) was used as the reference genome for all *L. monocytogenes* comparisons and annotation. Genomes of *essC8 L. monocytogenes* were downloaded from NCBI and reannotated using Prokka (v1.14.6) via the Galaxy server with default settings (35, 36) (https://usegalaxy.eu/). The presence of prophages in genomes was determined through submission to PHAge Search Tool Enhanced Release (PHASTER) web server (37, 38) (https://phaster.ca/).

### Identification of genetic hotspots and T7SS-related genes

Genomes of *essC8* variant *L. monocytogenes* genomes were directly aligned against *L. monocytogenes* EGDe (*essC1*) using progressiveMauve (39) (https://darlinglab.org/mauve/mauve.html). Genes within previously identified genetic hotspot regions were extracted, with protein accessions obtained via Batch Entrez, and homologous proteins identified from the NCBI reference proteins (refseq_protein) database for *L. monocytogenes* (taxid: 1639) via Blast under default search settings. Identification of EssC in non-*monocytogenes* species was carried out with blastp under the same conditions.

### Sequence alignment and protein domain prediction

Proteins of interest for this analysis were aligned using MUSCLE (40, 41) (https://www.ebi.ac.uk/Tools/msa/muscle/) to determine sequence identity with previously predicted T7SSb proteins. Protein alignment of conserved residues were visualised with Plotcon, with a window size of 5 (https://www.bioinformatics.nl/cgi-bin/emboss/plotcon). Potential toxin domains were predicted through the submission of protein sequences to Motif Search, against the Pfam database, with default settings (https://www.genome.jp/tools/motif/). Protein structural features were predicted either through SignalP 5.0 for the presence of secretory signal peptides (42) (https://services.healthtech.dtu.dk/service.php?SignalP-5.0) or through DeepTMHMM for the presence of transmembrane regions (43) (https://dtu.biolib.com/DeepTMHMM). Computational models of protein structures were obtained using AlphaFold (v2.2.4) colab (44) (https://colab.research.google.com/github/deepmind/alphafold/blob/main/notebooks/AlphaFold.ipynb). Prediction of biological activity for proteins of interest was via submission of computational models to the DALI server and PHYRE2 against known structures in the Protein Data Bank (45) (http://ekhidna2.biocenter.helsinki.fi/dali/).

### Phylogenetic analysis

Amino acid sequences of EssC variants from *L. monocytogenes* and non-*monocytogenes Listeria* spp. were aligned using MUSCLE. Phylogenetic trees were constructed using MEGA7, using the Neighbour-Joining algorithm, JTT matrix and pairwise deletions, with 100 bootstraps (46).

### Assessment of flanking genes

To identify conserved flanking genes in genetic hotspots between both *L. monocytogenes* and non-monocytogenes *Listeria* genomes, protein accessions were submitted to webFlaGs (47) (http://www.webflags.se/). Conserved genes were extracted, with genetic neighbourhood differences visualised with Clinker (48) (https://github.com/gamcil/clinker).

## RESULTS

### Identification of a novel variant of *essC* in Lineage III genomes of *L. monocytogenes*

Previously we had analysed the genetic arrangement of the T7SSb gene cluster across 271 *L. monocytogenes* genome sequences that were present in the NCBI database, resulting in the identification of seven sequence variants of EssC (22). In this study, we have expanded this analysis to investigate the distribution of *L. monocytogenes essC* variants present in a genome repository that was assembled from multiple sources, comprising 37,930 genomes that encompassed all four evolutionary lineages (I – IV). Within the repository there were 17,687 sequences from Lineage I genomes, 19,034 from Lineage II genomes, 1,133 from Lineage III genomes and 76 from Lineage IV genomes. These had been isolated from a diverse range of geographical locations and time periods, and from human, animal, food, factory and environmental sources (Table S1).

Genomes were phylogenetically clustered into their evolutionary lineages, annotated with their respective clonal complexes, and annotated as clinical or non-clinical (Fig. S1). The EssC-coding sequence was extracted from every genome and aligned against representative EssC1 – EssC7 sequences to classify the T7SSb subtype of each strain. The distribution of *essC* genetic variants across these evolutionary lineages and clonal complexes is summarised in Fig. 1a, with the distribution of *essC* genetic variants across all genomes in the repository shown in Fig. S1. Within Lineages I and II, there were clear associations of *essC* variant with the MLST clonal complexes, for example in Lineage I where CC1 exclusively encodes *essC1*, but CC2 and CC6 encode only *essC2*; similarly, in Lineage II, CC9 encodes *essC1*, but CC8 encodes *essC2*. The CC14 clonal complex in Lineage II is split into three distinct groups, respectively, and this shows in their respective *essC* variants. The first two clusters of CC14 encode *essC2* and *essC1* (Fig. S1), while the third cluster of CC14 encodes *essC1* and *essC5*. In contrast, in Lineages III and IV there is much more diversity in *essC* types, which is likely related to the much higher genomic variability in these lineages, as well as them being less often isolated and hence less represented in the repository. While there were clear associations with MLST clonal complexes, there was no link between *essC* variant and whether the isolate was from a clinical or non-clinical source.

**Figure 1.**
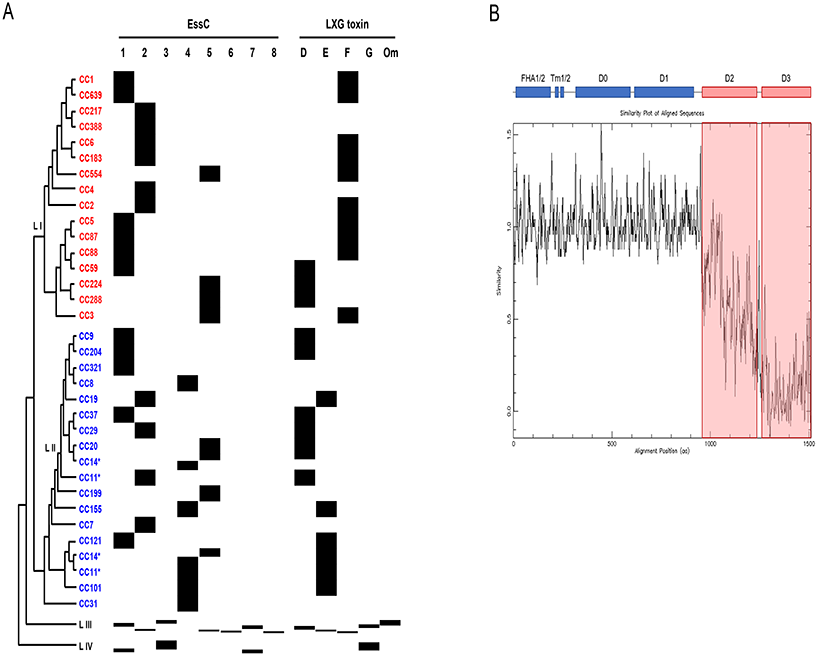
Schematic representation of the distribution of the eight *essC* genetic variants across *L. monocytogenes* lineages and clonal complexes. A. Tree of the 16 most common clonal complexes of *L. monocytogenes* lineages I and II, based on the order in Fig. S1 and Table S1. The tree is not representative of the number of isolates per clonal complex. EssC subtype and presence/identity of LXG toxin in variable region 2 is shown. Om – omicron. B. Plotcon analysis showing diversity across the eight EssC sequences from representative genomes, the domain organisation of EssC is shown above. FHA – forkhead associated domain; TM – transmembrane domain; D0, D1, D2 and D3 - ATPase domains.

Two of the 37,930 genome sequences did not contain an *essC* gene, however on further inspection these genome sizes (2.512Mb and 2.808Mb) were smaller than the 2.944Mb EGDe genome, and additionally lacked numerous essential genes as well as the T7SSb locus, so likely represent incomplete sequences. Of the remaining genomes, the *essC1* variant is the major subtype, with *essC2* the second most abundant (Table 1), in agreement with previous analysis of a much smaller number of genomes (22). We also noted previously that some *essC1* genomes harboured a second, truncated *essC* pseudogene encoding a protein from residue 771 onwards, but with a variable region from a different EssC. This finding was further reinforced in our larger dataset, and we noted that some *essC2* and *essC3* genomes also encoded a second truncated variant (Table 1). As reported previously (22), where a second partial *essC* copy was present, these genomes encode the predicted toxins of both *essC* variant types and contain a significantly larger variable region 1 compared to single copy *essC* genomes (22).

**Table 1.** Distribution of *essC* sequences across the *L. monocytogenes* genome repository. *Manual inspection of genomic sequences of these two genomes identified a sequencing gap covering the T7SS locus and multiple essential genes and therefore likely represent incomplete genomes.

| Encoded variant | Number of genomes | Overall percentage (%) |
| --- | --- | --- |
| EssC1 | 17063 | 44.985 |
| EssC2 | 9738 | 25.674 |
| EssC3 | 518 | 1.366 |
| EssC4 | 5755 | 15.173 |
| EssC5 | 4170 | 10.994 |
| EssC6 | 57 | 0.150 |
| EssC7 | 346 | 0.912 |
| EssC8 | 10 | 0.026 |
| Full-length EssC1/truncated EssC2 | 104 | 0.274 |
| Full-length EssC1/truncated EssC3 | 126 | 0.332 |
| Full-length EssC1/truncated EssC4 | 33 | 0.087 |
| Full-length EssC1/truncated EssC5 | 2 | 0.005 |
| Full-length EssC2/truncated EssC3 | 1 | 0.003 |
| Full-length EssC3/truncated EssC4 | 3 | 0.008 |
| Full-length EssC3/truncated EssC5 | 1 | 0.003 |
| Full-length EssC3/truncated EssC7 | 1 | 0.003 |
| No EssC * | 2 | 0.005 |
| Total | 37930 | 100 |

During our analysis we noted that EssC sequences from a small subset of ten Lineage III genomes clustered separately from the seven EssC variants we had identified previously, forming an eighth EssC variant cluster. All ten of these *essC8* genomes were isolated in the USA, and no one source category could be assigned to all genomes. These ten genomes are listed in Table 2, alongside metadata regarding isolation details, and a sequence alignment of EssC8 against the other seven EssC variants is shown in Fig. S2. Ultimately, this rare variant accounted for 0.88% of genomes in Lineage III of *L. monocytogenes*, and 0.026% of all genomes examined in this analysis. Following this, we later identified a further *essC8*-encoding strain - the recently sequenced soil isolate *L. monocytogenes* FSL L7-0325 - after BLAST analysis of the NCBI RefSeq database, which we also included in our subsequent bioinformatic analysis.

**Table 2.** *L. monocytogenes essC8* genomes from the *L. monocytogenes* genome repository.

| Strain | Lineage | Clonal complex | Source category | Collection date | Location | Biosample | Assembly |
| --- | --- | --- | --- | --- | --- | --- | --- |
| 2017L-6133 | III | ST1636 | Human | No year | USA | SAMN11320611 | GCA_004764935.1 |
| PNUSAL002887 | III | ST1636 | Human | 2017 | USA | SAMN07503874 | GCA_004607655.1 |
| AR-FR-3-WH1 | III | N/A | Environment | 2018 | USA | SAMN12630146 | GCA_008270535.1 |
| PNUSAL006195 | III | N/A | No source | 2019 | USA | SAMN13981699 | GCA_010354525.1 |
| PNUSAL006196 | III | N/A | No source | 2019 | USA | SAMN13981723 | GCA_010357625.1 |
| 13FMFO000101 | III | N/A | Food | 2013 | USA | SAMN06899320 | GCA_004724445.1 |
| CFSAN061343 | III | ST1358 | Animal | 2014 | USA | SAMN17814432 | GCA_017122535.1 |
| PNUSAL004142 | III | ST2005 | Human | 2018 | USA | SAMN09761446 | GCA_003678025.1 |
| PNUSAL004951 | III | ST2065 | Food | 2019 | USA | SAMN12142118 | GCA_006495195.1 |
| PNUSAL004950 | III | ST2065 | Food | 2019 | USA | SAMN12142119 | GCA_006494955.1 |

In addition to EssC, the T7SSb of Bacillota comprises three additional membrane proteins, EsaA, EssA and EssB, and two globular components, EsaB and EsxA (49). Our previous analysis had indicated that across the *L. monocytogenes essC1-essC7* genomes, these components are highly conserved (approximately 97% identity, (22)). Extending our analysis to *essC8* genomes we noted that these five core proteins were also highly similar to those encoded by the other seven variant genomes (e.g. EsxA ≥96.6%, EsaB ≥98.8%). By contrast, the sequence identity of EssC8 relative to the other seven EssC variants ranged from 77.53% with EssC4 to 82.80% with EssC1, with the variability almost exclusively encompassing the cytosolic D2 and D3 portions of the protein (Fig. S2). Submission of representative EssC sequences of each variant to PlotCon confirmed that sequence variation at the protein level is confined mainly to the C-termini of these proteins (Fig. 1B).

### *essC8* variable region 1 encodes an Rhs toxin

The T7SSb locus of *L. monocytogenes* genomes comprises two variable regions that are separated by a cluster of five housekeeping genes (encoding predicted phosphoenolpyruvate mutase, 6-O-methylguanine DNA methyltransferase, a YjbI-superfamily protein, a 2-hydroxyacid dehydrogenase and a tRNA, respectively (Fig 2; (22)). Variable region 1 encodes one or more candidate substrate proteins that are specific for a particular *essC* variant; these include a YeeF-like domain toxin in *essC1* genomes and LXG toxins A, B and C in *essC5*, *essC6* and *essC7* genomes, respectively (22).

**Figure 2.**
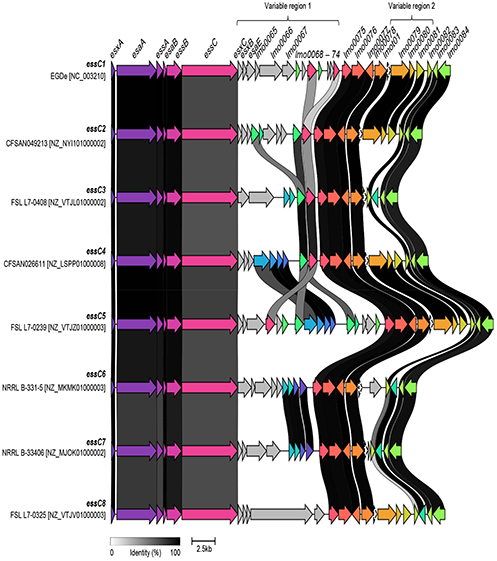
The T7SS locus of representative examples of each *L. monocytogenes essC* variant. Variable regions 1 and 2 are shown. Genes shown in grey in variable region 1 are *essC*-subtype specific; otherwise homologous genes are shaded in the same colour and the black/grey blocks indicate the level of identity across homologues.

Fig. 2 provides an overview of the T7SSb locus of an example of each *essC* variant, with homologous genes conserved across the eight genomes shaded by sequence similarity. As expected, variable region 1 is highly heterogeneous between the eight *essC* subtypes, since each *essC* strain encodes variant-specific substrate proteins (usually alongside cognate immunity proteins). However, we noted previously that genes encoding ‘orphan’ immunity proteins (i.e. to protect from ‘non-self’ toxins) are also encoded in variable numbers in almost all genomes (22). In the particular example in Fig. 2, the predicted cognate immunity gene to the LXG-domain toxin of the *essC5* strain FSL L7-0239, is also encoded in variable region 1 of the *essC1*, *C2*, *C3* and *C4* genomes.

The dominant genetic feature of variable region 1 for all *essC8* genomes is the presence of a large gene encoding a 1656-residue protein (Fig. 2, Fig. 3a, Fig. 4). The protein has a DUF6531 (PF20148) domain that precedes 16 rearrangement hotspot (Rhs) repeat (PF05593) domains extending down its length (Fig. 3b). We subsequently renamed this protein *Listeria* Rhs toxin A (LrhA). Many Rhs-repeat proteins are anti-bacterial toxins that can be secreted via a number of routes, including the Gram-negative Type VI secretion system and by the Sec pathway in Gram-positive bacteria, dependent upon the targeting information that is present (50, 51). The Rhs repeats form a filamentous ‘cocoon’ that encases a C-terminal toxin domain (52).

**Figure 3.**
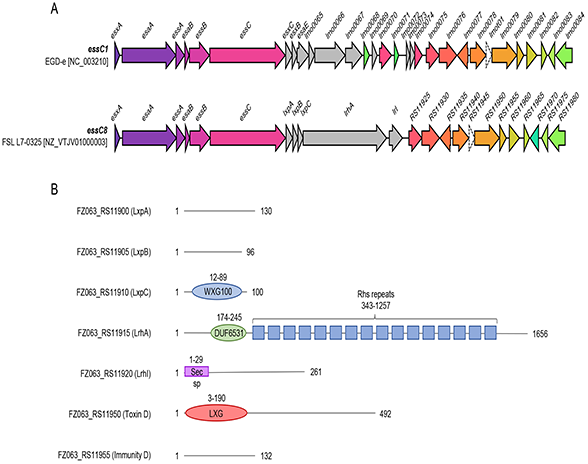
*L. monocytogenes essC8* genomes encode an Rhs toxin. A. Schematic representation of the T7SS loci of the EGDe type strain (*essC1* variant) with FSL L7-0325 (*essC8* variant). B. Domain predictions for proteins encoded at the FSL L7-0325 *essC8* locus. sp – signal peptide.

**Figure 4.**
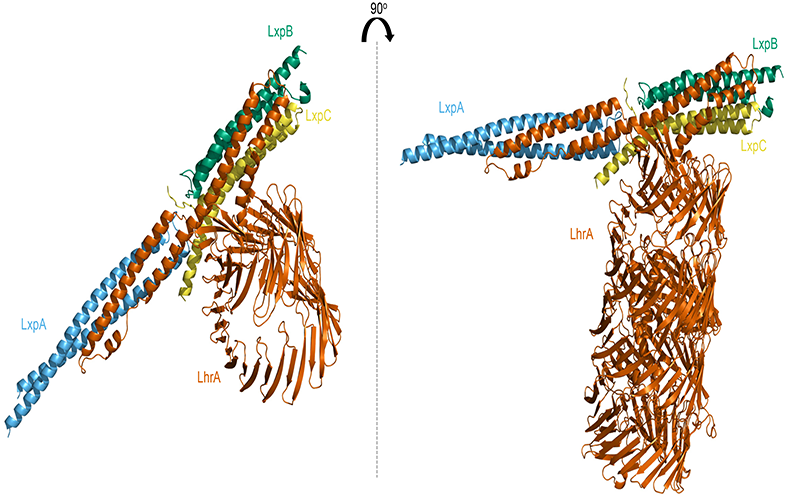
Predicted structures of the Rhs protein LrhA in complex with likely interacting partner proteins LxpABC.

Immediately downstream of *lrhA* is a candidate immunity gene which we have termed *lrhI*. The *lrhI* gene encodes a 261-residue protein with a predicted transmembrane domain close to its N-terminus; no protein family could be assigned to the remaining portion of this protein. As the globular portion of LrhI is predicted to be extracellular, the target of LrhA is likely to be within the cell envelope. Consistent with our annotation of LrhI as an immunity protein, ‘orphan’ copies are encoded in variable region 1 of multiple other *L. monocytogenes essC* types, including genomes UTK_C1-0018-E1 (*essC1*; *LAX90_RS06725*), UTK_C1-0003-E1 (*essC2*; *LAX77_RS05315*) and NRRL B-3381 (*essC5*; *BJM68_RS12710*).

Most T7SSb toxins analysed to date are encoded at loci that also contain genes for small helical proteins, some of which are from the WXG100 family and that serve as targeting factors/chaperones for toxin secretion (16, 19). Sandwiched between *essC8* and *lrhA* are three small genes. One of these (FSL L7-0325, *FTZ063_RS11910*, Fig. 3b) encodes a canonical WXG100 family protein, while the other two also encode small proteins that are predicted to be almost completely alpha-helical.

### Modelling of LrhA confirms predicted Rhs structural features

To explore the organisation of LrhA further, including identifying any potential structural features of the N-terminal region (which has no predicted homology to any known protein domain), we divided the protein sequence into three equal sections with 100 residue overlaps and analysed them with AlphaFold multimer, with each output being aligned to generate a model of the entire protein. The AlphaFold prediction clearly modelled the Rhs repeats as a column of β-sheets, with the folded C-terminus oriented within the β-sheet bundle (Fig. 4). While AlphaFold also predicted the fold of the C-terminal toxin domain, we were not able to identify any significant structural matches to this protein region using the DALI or PHYRE2 servers.

To explore any potential protein-protein interactions, the three small proteins encoded upstream of *lrhA* were modelled alongside the LrhA N-terminal 173 amino acids using AlphaFold multimer. This resulted in all three proteins being positioned at the α-helical N-terminus of LrhA, with the α-helices aligned along a single axis, almost intercalating with one another. Based on this model, together within the current understanding in the role of these small proteins in T7 substrate secretion in other species (16, 19), these proteins were assigned the term *Listeria* T7SSb auxillary proteins (LxpABC). The predicted AlphaFold model of LrhA bound to the Lxp proteins is shown in Fig. 4.

### A novel LXG-domain toxin, Omicron, is encoded at variable region 2 of some genomes

The second variable region found at the *L. monocytogenes* T7SSb locus, variable region 2, is bounded by a tRNA-lysine gene, *lmot01* at the 5′ end and a hypothetical membrane protein-encoding gene, *lmo0082* at the 3′ end. This variable region is generally much shorter than variable region 1, encoding a single LXG-domain toxin (either toxin D, E, F or G; (22)) alongside a cognate immunity gene and sometimes additional ‘orphan’ immunities. We noted that in the small dataset analysed previously, approximately 25% of genomes had no toxin encoded at this locus (22).

Analysis of our larger dataset revealed that 74.6% of the genomes encoded an LXG-domain toxin in variable region 2 (Table 3, Tables S2-S5, Fig. 1A, Fig. S1); we never found more than one LXG toxin encoded in this region. This heterogeneity was also borne out in the 11 *essC8* genomes where seven of them were found to encode an LXG-domain protein. Across these seven genomes, one of three LXG-domain toxins were present. Three genomes encoded a copy of Toxin G, while two encoded copies of Toxin D. Surprisingly, two genomes encoded a previously unidentified toxin at this position, designated Toxin Omicron (Table 3). An alignment of Omicron with Toxins D – G (Fig. S3a) and plotcon analysis (Fig S3b) shows that it shares the same highly conserved N-terminal region (up to amino acid 343), with a divergent C-terminal toxin domain that shares structural similarity to the Tne2 NADase Type VI secretion system effector (53). A predicted immunity gene for Omicron is encoded adjacent to the toxin and is also found as an orphan immunity gene in variable region 2 of some other *L. monocytogenes* genomes.

**Table 3.** Occurrence of five different LXG-domain toxins in variable region 2 of genomes from the *L. monocytogenes* genome repository.

| EssC variant | LXG N-terminus detected | Toxin D | Toxin E | Toxin F | Toxin G | Toxin Omicron | No LXG toxin | Total |
| --- | --- | --- | --- | --- | --- | --- | --- | --- |
| EssC1 | 14688 | 4283 | 2022 | 8334 | 32 | 14 | 2375 | 17063 |
| EssC1/EssC2 (truncated) | 19 | 19 | 0 | 0 | 0 | 0 | 85 | 104 |
| EssC1/EssC3 (truncated) | 27 | 27 | 2 | 0 | 13 | 9 | 99 | 126 |
| EssC1/EssC4 (truncated) | 32 | 32 | 1 | 0 | 0 | 0 | 1 | 33 |
| EssC1/EssC5 (truncated) | 2 | 2 | 0 | 1 | 0 | 0 | 0 | 2 |
| EssC2 | 6141 | 6141 | 674 | 4457 | 8 | 12 | 3597 | 9738 |
| EssC2/EssC3 (truncated) | 1 | 1 | 0 | 0 | 0 | 1 | 0 | 1 |
| EssC3 | 216 | 216 | 3 | 3 | 66 | 74 | 302 | 518 |
| EssC3/EssC4 (truncated) | 0 | 0 | 0 | 0 | 0 | 0 | 3 | 3 |
| EssC3/EssC5 (truncated) | 0 | 0 | 0 | 0 | 0 | 0 | 1 | 1 |
| EssC3/EssC7 (truncated) | 1 | 1 | 0 | 0 | 1 | 0 | 0 | 1 |
| EssC4 | 3383 | 3383 | 3322 | 5 | 2 | 0 | 2372 | 5755 |
| EssC5 | 3505 | 3505 | 310 | 1911 | 15 | 7 | 665 | 4170 |
| EssC6 | 28 | 28 | 0 | 0 | 8 | 2 | 29 | 57 |
| EssC7 | 251 | 251 | 0 | 13 | 24 | 210 | 95 | 346 |
| EssC8 | 6 | 6 | 0 | 0 | 3 | 2 | 4 | 10 |
| Total | 28300 | 6731 | 6334 | 14724 | 172 | 331 | 9628 | 37928 |

Although we had not identified Toxin Omicron in our previous analysis, Omicron was also encoded at low frequency in variable region 2 of other genomes in our larger dataset (Table 3). Further analysis revealed that all of these are exclusively Lineage III genomes (Table S4). This prompted us to examine whether other variable region 2 toxins showed any lineage specificity. Strikingly, it is clear that LXG toxin identity clustered with EssC type across lineage and MLST clonal complex, but that this was irrespective of whether strains were isolated from the environment or clinical settings (Fig. 1A, Fig. S1). We noted that only Toxins D or F were encoded across Lineage I (with Toxin F being the most frequent; Table S2, Fig. 1A) whereas Lineage II genomes predominantly encode Toxin D or E, with rare instances of Toxin F (Table S3, Fig. 1A). Lineage III genomes showed the highest variability, with instances of genomes encoding each of the five different toxins. By contrast, the small number of Lineage IV genomes in our dataset only encoded Toxin H, and the majority of genomes in this lineage did not encode any LXG toxin at this region (Table S5, Fig. 1A).

### Insertion of a prophage within variable region 2 of *essC8* strain 13FMFO000101

During examination of the *essC8* genomes it was noted that *L. monocytogenes* 13FMFO000101 presented an extensively extended variable region 2 due to insertion of 38.6kb prophage into the tRNA gene, *lmot01* (Fig. 5). While no direct identity could be assigned to the prophage, it was determined to be intact by the phage prediction PHASTER webserver, with 64 ORFs being identified. Interestingly, insertion of the prophage did not disrupt the LXG-domain toxin or immunity genes in this region. Furthermore, the *attR* arm of the prophage did not disrupt the tRNA-Lysine as a result of insertion into the chromosome. None of the prophage ORFs were predicted to encode any T7SSb-related toxin or immunity genes. An overview of the ten *essC8* genomes identified from the genome repository is illustrated in Fig. 5, including the highly extended variable region 2 of 13FMFO000101.

**Figure 5.**
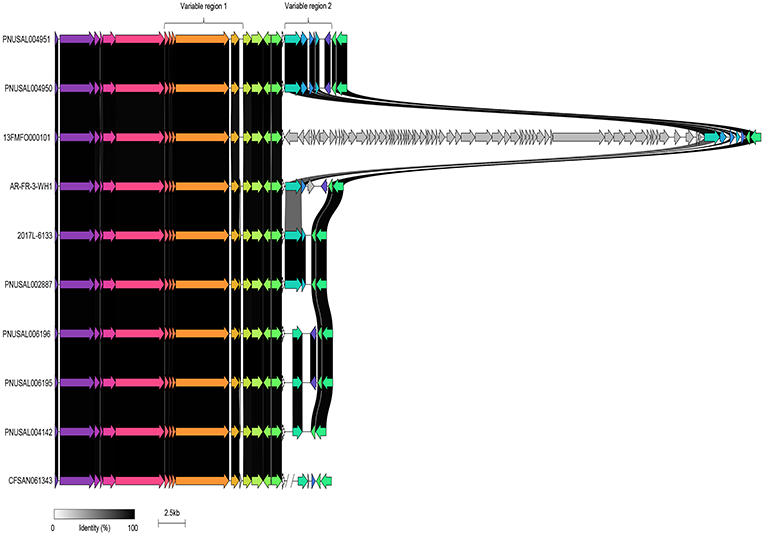
The *essC8* strain 13FMFO000101 has a prophage insertion in variable region 2. The *ess* locus of 13FMFO000101 alongside the further nine *essC8 L. monocytogenes* genomes identified in the repository is shown. Homologous genes are shaded in the same colour and the black/grey blocks indicate the level of identity across homologues.

### Identification of new *ess*-external hotspots and predicted T7SSb substrates

Previously, we identified ten chromosomal hotspots across *L. monocytogenes*, in addition to variable regions 1 and 2, where LXG-domain substrates could be encoded (Fig. 6), and 33 distinct LXG toxin sequences (Toxins H – ϕ, Table S6) (22). When a hotspot was ‘occupied’, generally only one LXG-domain-coding gene was found (except for the hotspot at *lmo1096* where two LXG proteins were encoded in a head-to-head arrangement). Additional features of hotspots are the presence of one or two genes encoding small helical hairpin proteins sharing the WXG100 family fold, along with genes for immunity proteins (22).

**Figure 6.**
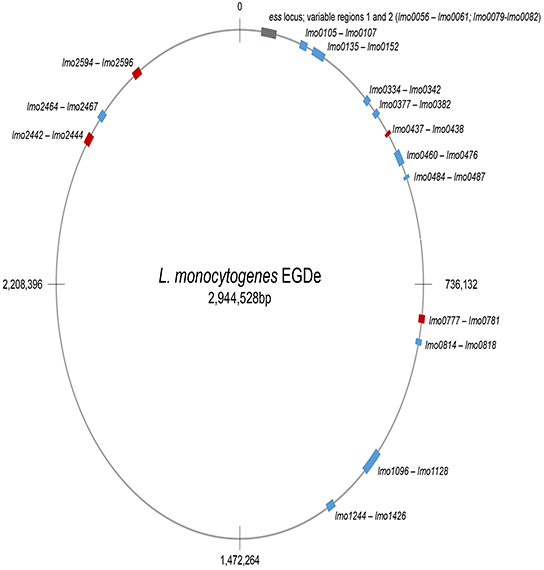
Chromosomal ‘hotspots’ where LXG-protein-encoding genes occur in *L. monocytogenes* genomes. Hotspots shown in blue have been identified previously (22), those in red were found following genome analysis of *essC8* genomes.

To examine the repertoire of LXG-domain proteins in *essC8* genomes, we searched genome annotations with the term ‘LXG’, and also undertook BLAST analysis of encoded proteins using the LXG domains of previously-identified *L. monocytogenes* toxins (22) (Table S6). From this we found that the *essC8* genomes collectively encoded a further seventeen of the LXG proteins we had described previously (22), in addition to those at variable region 2. We also noted that as well as Omicron, there were a further eight LXG-domain proteins that were novel. In each case, the LXG proteins were encoded adjacent to a probable immunity gene and were preceded by one or two genes for WXG100-like proteins. Gene neighbourhood analysis showed that four of these novel toxins - Nu, Ψ, Iota and ω - were encoded at known hotspot regions (Table S6). The other four LXG-proteins – Kappa, ξ, Upsilon and Chi were encoded at four new hotspots (Fig 6, Table S6). BLAST searches indicated that all eight of these toxins were encoded, at varying frequencies, in other *essC* variant genomes.

### The T7SSb is encoded by non-monocytogenes *Listeria* spp

A blastp search identified that EssC homologues are encoded in many species of *Listeria* in addition to *L. monocytogenes*. Batch Entrez was used to retrieve available EssC sequences of non-monocytogenes *Listeria* species from the RefSeq database and an alignment of these sequences was performed, with a representative of each *L. monocytogenes* EssC (1–8) for reference. A Neighbour-Joining tree is shown in Fig. 7. All eight EssC variants are represented in non-*monocytogenes Listeria* spp., with *L. innocua*, *L. welshimeri* and *L. seeligeri* encoding seven of the eight EssC variants including EssC8 (Table 4). Outside of EssC 1-8, other potential EssC types were detected, including four possible types primarily associated with *L. booriae*, three possible types with *L. grayi*, and six other possible types with other *Listeria* species (Fig. 7, Table 4).

**Table 4.** Distribution of EssC1-8 and novel EssC variants in the genus *Listeria*. *Based on distance in phylogenetic tree comparing amino acid sequences (Fig. 7).

| **EssC variant** | **Present in *Listeria* species:** |
| --- | --- |
| EssC1 | *L. monocytogenes*, *L. innocua*, *L. marthii*, *L. welshimeri*, *L. seeligeri*, *L. ivanovii* |
| EssC2 | *L. monocytogenes*, *L. innocua*, *L. cossartiae*, *L. seeligeri* |
| EssC3 | *L. monocytogenes*, *L. innocua*, *L. marthii*, *L. cossartiae*, *L. farberii*, *L. welshimeri*, *L. ivanovii*, *L. immobilis*, *L. seeligeri* |
| EssC4 | *L. monocytogenes*, *L. innocua*, *L. marthii*, *L. cossartiae*, *L. welshimeri* |
| EssC5 | *L. monocytogenes*, *L. innocua*, *L. welshimeri*, *L. seeligeri*, *L. ivanovii* |
| EssC6 | *L. monocytogenes*, *L. marthii*, *L. cossartiae*, *L. welshimeri*, *L. seeligeri* |
| EssC7 | *L. monocytogenes*, *L. innocua*, *L. marthii*, *L. cossartiae*, *L. seeligeri*, *L. immobilis*, *L. welshimeri* |
| EssC8 | *L. monocytogenes*, *L. swaminathanii*, *L. seeligeri*, *L. welshimeri* |
| Novel* | *L. costaricensis* |
| Novel* | *L. ilorinensis* |
| Novel* | *L. valentina*, *L. kieliensis*, *L. aquatica* |
| Novel* | *L. fleischmannii*, *L. goaensis* |
| Novel* | *L. floridensis* |
| Novel* | *L. booriae*, *L. weihenstephanensis*, *L. rustica*, *L. newyorkensis*, *L. rocourtiae* |
| Novel* | *L. booriae*, *L. grandensis* |
| Novel* | *L. booriae*, *L. grandensis* |
| Novel* | *L. grayi* |
| Novel* | *L. grayi* |
| Novel* | *L. grayi* |

**Figure 7.**
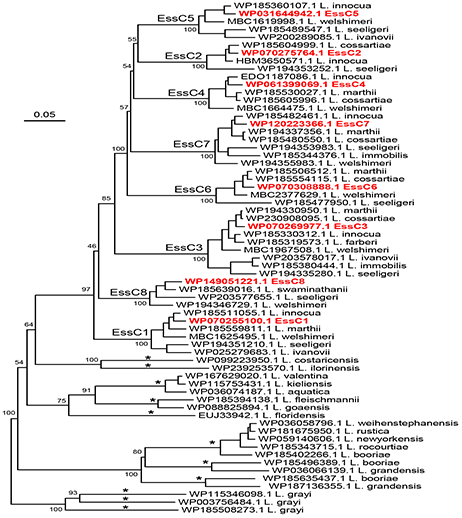
Phylogenetic tree of selected EssC sequences from across the *Listeria* genus, based on an alignment of predicted amino acid sequences. Examples of *L. monocytogenes* EssC1 – EssC8 sequences were included in the analysis. The names represent the protein accession number with either the Listeria species of *L. monocytogenes* EssC variant. Asterisks on tree branches indicate 11 possible new EssC variants in non-monocytogenes *Listeria* species (see also Table 4). Values at the major branching nodes represent bootstrap values, based on 100 iterations.

### The T7SSb Rhs toxin, LrhA, is encoded by *L. welshimeri*, *L. seeligeri* and *L. swaminathanii*

To determine if the Rhs toxin, LrhA was restricted to *L. monocytogenes* or more broadly distributed in the *Listeria* genus, a blastp search was performed. We used Batch Entrez to download available accession numbers from RefSeq, and analysed these by gene neighbourhood analysis (Fig S4). From this it was apparent that LrhA homologues are also found in *L. welshimeri, L. swaminathanii* and *L. seeligeri* (Fig. S4). Interestingly, while in *L. monocytogenes, L. swaminathanii* and *L. welshimeri* LrhA is only encoded downstream of *essC8*, in *L. seeligeri* LrhA could additionally be found in association with EssC2, EssC5 and EssC7 (Fig. 8). As discussed above, a small fraction of *L. monocytogenes* genomes code for a second, truncated copy of EssC of a different variant. We noted that some of the *L. seeligeri* genomes also have truncated *essC* genes and that LrhA was also encoded downstream of truncated *essC2*, *essC5*, *essC7* and *essC8* (Fig. 8).

**Figure 8.**
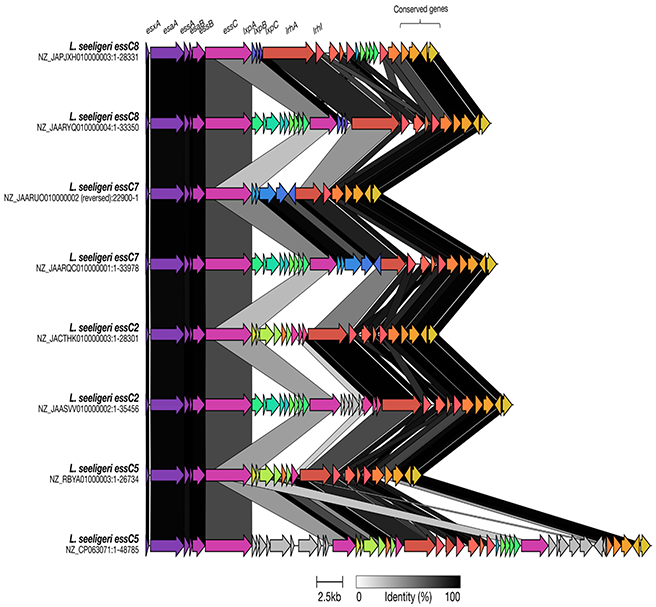
Genomic organisation of *lrhA* genes in *L. seeligeri* species. A representative truncated LrhA homologue was taken from each *essC* variant identified in *L. seeligeri*. Homologous genes are shaded in the same colour and the black/grey blocks indicate the level of identity across homologues.

When we compared the sequence of LrhA encoded downstream of the different EssC variants we noticed that the EssC2, EssC5 and EssC7-associated LrhAs were shorter than the EssC8-associated protein (Fig. S5). The truncated LrhAs in these genomes lack the helical N-terminal region and DUF6531 domain, comprising only Rhs repeats and the C-terminal toxin domain. They also lack the three genes encoding the LxpABC partner proteins. It is not clear whether any of these truncated LrhA proteins would be substrates of the T7SSb, and if so how they would be targeted to the secretion system. We also note that although the Rhs repeat sequence is highly conserved, the toxin domain is quite divergent across these four LrhA variants. Rhs protein-encoding genes are notorious sites for recombination (e.g. (54)), and it is possible that these truncated *lrhA* genes act as a source of sequence variability for the *L. seeligeri* species, providing a template for diversification of the toxin domain to overcome immunity.

## DISCUSSION

In this work we have assembled a repository of 37,930 *L. monocytogenes* genomes and interrogated it to explore the diversity in the T7SSb and its substrates across the species. We have shown that the core secretion component, EssC, occurs as one of eight variants, with EssC1 being the most common and EssC8 the rarest. We also show that a small subset of *essC1*, *essC2* and *essC3* genomes encode a truncated EssC of a different variant, along with the likely substrate protein/s recognised by that specific EssC type. Four variants of EssC have previously been reported in *S. aureus* (55), and recently it was shown that there are four EssC types in group B *Streptococcus* (56), suggesting that heterogeneity is a general feature of the T7SSb.

The EssC8 variant had not been previously described and at present is only found in a few *L. monocytogenes* Lineage III genomes. An Rhs domain protein, LrhA, is encoded downstream of *essC8* but is not found in other *essC* variant genomes and we suggest that in *L. monocytogenes* it is a T7SSb-EssC8-specific substrate. This is further supported by the finding that LrhA is encoded adjacent to genes for small WXG100 proteins, which are known to act as chaperones and/or targeting factors for T7-secreted toxins (16, 19).

The most common toxin substrates of the T7SSb harbour an LXG domain and our prior analysis had identified 40 distinct LXG proteins across *L. monocytogenes* genomes (22). While we were not able to individually analyse all of the genomes in the repository, inspection of the *essC8* genomes identified a further nine proteins of the LXG family. Some of these were encoded within known chromosomal hotspots, whereas four were encoded at new hotspots not previously described. Retrospective analysis indicated that these novel LXG proteins were not unique to *essC8* genomes but also present in other *essC* variant genomes in the repository. It is likely that there are additional LXG proteins encoded within the species that may be revealed by more intensive study of the available genome sequences.

We extended our analysis to assess T7SSb variability in the *Listeria* genus. Although there is significantly less genome sequence information outside of *L. monocytogenes*, we were able to identify numerous subtypes of EssC in other *Listeria* spp., including examples of each of the eight *L. monocytogenes* EssC variants. We also identified several novel EssC variants, and for species where multiple complete genome sequences were available, we frequently found they encoded more than one EssC subtype. Taken together this indicates that the genetic diversity we have reported for *L. monocytogenes* T7SSb is likely to be a consistent feature across the genus.

The Rhs toxin identified in *L. monocytogenes essC8* genomes was also found in other species. While in *L. welshimeri* and *L. swaminathanii* LrhA was found solely in association with EssC8, in *L. seeligeri* it was also encoded in *essC2*, *essC5* and *essC7* genomes. Curiously, however, in these three *essC* variants LrhA was truncated, lacking the apparent T7SSb helical targeting domain at the N-terminus, and the three *lxp* genes were also absent. It remains to be determined whether these are functional toxins or remnants that are not expressed or secreted.

Rhs proteins are large antibacterial toxins with polymorphic C-terminal domains that mediate contact-dependent growth inhibition (57), and prior analysis had identified a link between Rhs proteins and some secretion pathways, including the T7SS (58). The T7SS is known to export small helical hairpin substrates as folded dimers, which have a diameter of approximately 25 Å (12, 59). The maximum size of the T7SS secretion pore is unclear as cryo EM structures of the mycobacterial T7SSa are of the closed complexes (7, 9), but modelling studies based on the related FtsK hexamer estimates a channel of approximately 30 Å (10). LrhA is the largest T7SSb substrate identified to date and is likely too large to be exported in a folded state; for example a typical Rhs cocoon such as that of *Pseudomonas protegens* RhsA is 86 x 65 Å (52). The Sec-dependent Rhs proteins would also be secreted unfolded and assume their 3D structure in the extracellular environment (51).

Analysis of the distribution of EssC variants and variable region 2 LXG toxin types showed clear associations with the respective genome lineage and multilocus sequence clonal complex. Within these clonal complexes there was relatively little, if any, variation in EssC and LXG toxin variants (Fig 1A, Figure S1), suggesting there is an effect of the genomic context of the T7SSb locus in restricting variation and genetic transfer within *L. monocytogenes*.

In conclusion, the extreme genetic variability in the *Listeria* T7SSb and its substrate proteins strongly suggests a major role for this system in bacterial antagonism. Further work is needed to determine under which conditions *Listeria* use T7SS-dependent competition.

## Author Contributions

K.B., T.P. and A.v.V. conceptualised the study. A.v.V. generated the genome repository of sequences. K.B. and A.v.V. analysed repository dataset. S.R.G. and A.v.V. analysed the non-monocytogenes dataset. K.B., S.R.G, A.v.V. and T.P. created the figures. K.B. and T.P. wrote the manuscript. All authors approved the final manuscript.

## Acknowledgements

We thank Emma Dobson for helping with preliminary bioinformatics work.

## Figure Legends

Figure S1. Phylogenetic trees of a) Lineage I b) Lineage II and c) Lineages III and IV of *L. monocytogenes* in the genome repository. The phylogenetic tree is based on core genome MLST profiles. The clonal complexes and clinical outcome are mapped onto each of the four linages, *essC* subtype and presence/identity of the LXG toxin in variable region 2 is also shown. Om – omicron.

Figure S2. Sequence alignment of the eight EssC variants found in *L. monocytogenes*. The shaded lines above the sequence indicate the approximate domain boundaries of EssC; red – FHA domains; grey – transmembrane domains; yellow – D0; blue – D1; orange – D2; green D3. The boxes indicate the percentage identity between each variant.

Figure S3. LXG toxins D, E, F, G and Omicron share a common N-terminal sequence. A. Sequence alignment of the five toxins. B. Analysis of the same five toxins using plotcon. The predicted boundary of the LXG domain is indicated.

Figure S4. Genomic organisation of *lrhA* genes in non-monocytogenes species of *Listeria*. WebFlags was used to generate a genetic neighbourhood analysis of all LrhA homologues found in non-monocytogenes species. The tree was generated by webFlaGs using the ETE3 toolkit.

Figure S5. Truncated variants of LrhA are encoded at the *ess* locus in *L. seeligeri* genomes. A. A representative of each truncated LrhA from the different *essC* variants was aligned using MUSCLE v3.8.1551 and visualised using Boxshade. B. Schematic representation the truncated LrhA proteins encoded by *L. seeligeri essC2*, *essC5* and *essC7* in comparison to full length LrhA. The red box at the beginning of the *essC7* Rhs variant denotes a stretch of 19 amino acids which do not directly align to the consensus sequence.

