## Supplemental Figures for "A novel variant of the *Listeria monocytogenes* type VII secretion system EssC component is associated with an Rhs toxin"

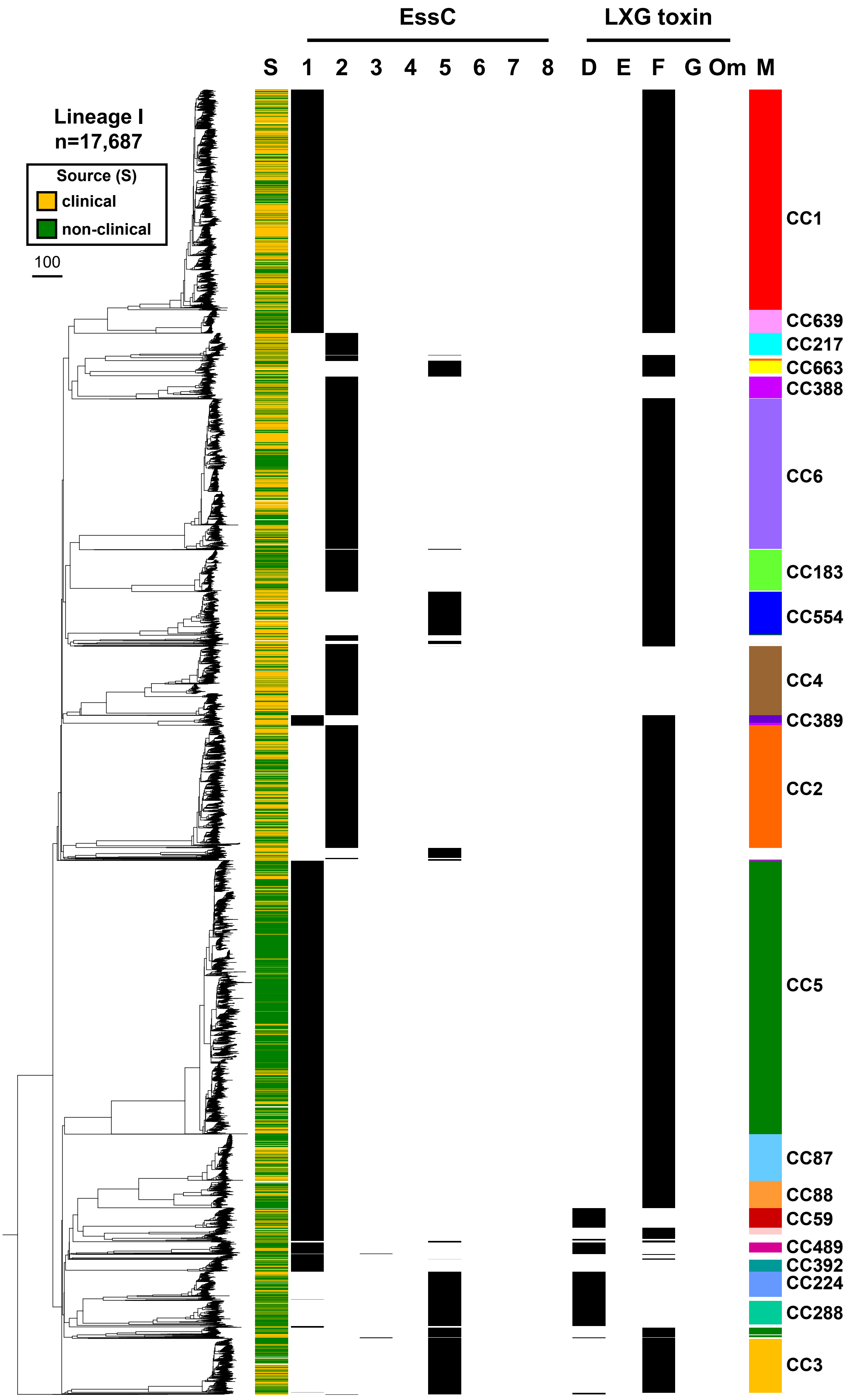

EssC
LXG toxin
S
1
2
3
4
5
6
7
8
D
E
F
G
Om
M
Lineage I
n=17,687
Source (S)
clinical
non-clinical
100
CC1
CC639
CC217
CC663
CC388
CC6
CC183
CC554
CC4
CC389
CC2
CC5
CC87
CC88
CC59
CC489
CC392
CC224
CC288
CC3

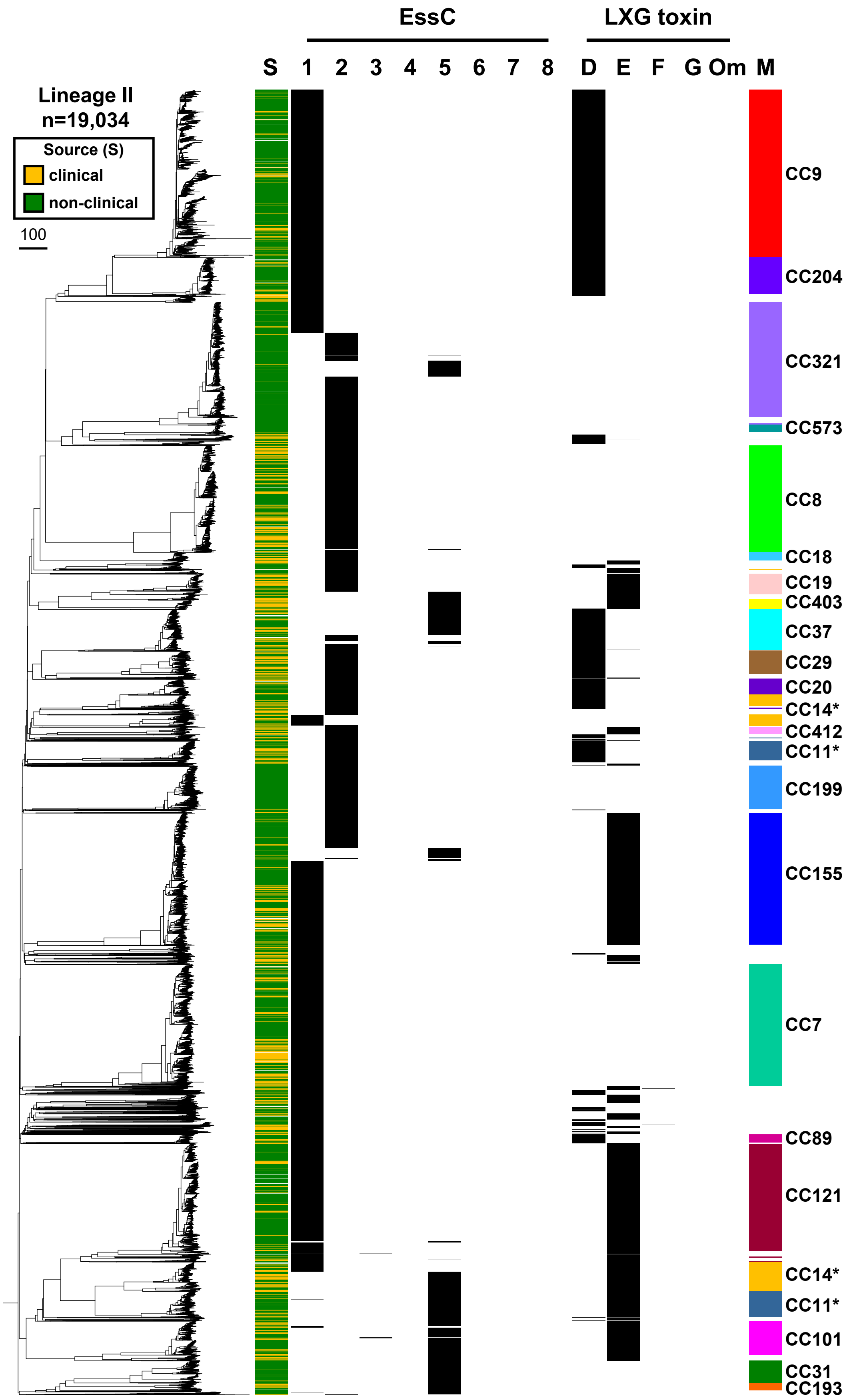

EssC
LXG toxin
S
1
2
3
4
5
6
7
8
D
E
F
G
Om
M
Lineage II
n=19,034
Source (S)
clinical
non-clinical
100
CC9
CC204
CC321
CC573
CC8
CC18
CC19
CC403
CC37
CC29
CC20
CC14*
CC412
CC11*
CC199
CC155
CC7
CC89
CC121
CC14*
CC11*
CC101
CC31
CC193

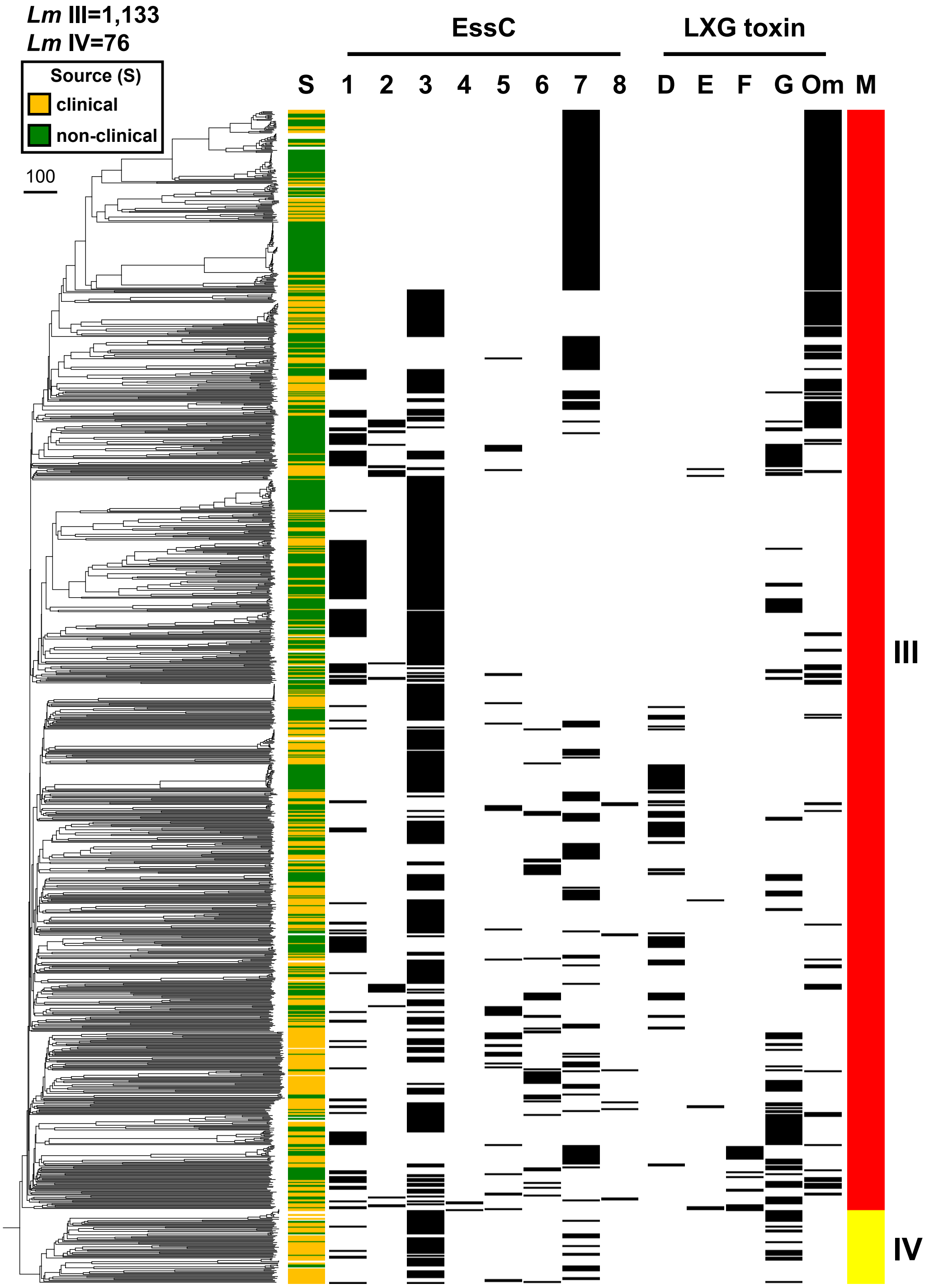

Figure S1. Phylogenetic trees of a) Lineage I b) Lineage II and c) Lineages III and IV of *L. monocytogenes* in the genome repository. The phylogenetic tree is based on core genome MLST profiles. The clonal complexes and clinical outcome are mapped onto each of the four linages, *essC* subtype and presence/identity of the LXG toxin in variable region 2 is also shown. Om – omicron.

```
EssC1   MNREALLIVSNGQQCHKNHLSPEKVVTIGNTIEHEITYPELAESIEVKYDEGSWNAGTTA  60
EssC2   MNREALLIVSNGQQCHKNHLSPEKVVTIGNTIEHEITYPELAESIEVKYDEGSWNAGTTA  60
EssC3   MNREALLIVSNGQQCHKHHLSPERVVTIGNTIEHEITYPELAESIEVKYDEGSWNAGTTA  60
EssC4   MNREALLIVSNGQQCHKNHLSPEKVVTIGNTIEHEITYPELAESIEVKYDEGSWNAGTTA  60
EssC5   MNREALLIVSNGQQCHKNHLSPEKVVTIGNTIEHEITYPELAESIEVKYDEGSWNAGTTA  60
EssC6   MNREALLIVSNGQQCHKHHLSPERVVTIGNTIEHEITYPELAESIEVKYDEGSWNAGTTA  60
EssC7   MNREALLIVSNGQQCHKHHLSPERVVTIGNTIEHEITYPELAESIEVKYDEGSWNAGTTA  60
EssC8   MNREALLIVSNGQQCHKHHLSPERVVTIGNTIEHEITYPELAESIEVIYDEGSWNAGTTA  60

EssC1   LQANQAVNVENLAFYLCQDLHTQVYDVVTNISVTFGVGIENDVTLDDTKSDFILLRDAKE  120
EssC2   LQANQAVNVENLAFYLCQDLHTQVYDVVTNISVTFGVGIENDVTLDDTKSDFILLRDAKE  120
EssC3   LQANQAVNVENLAFYLCQDLHTQVYDVVTNISVTFGVGIENDVTLDDTKSDFILLRDTKE  120
EssC4   LQANQAVNVENLAFYLCQDLHTQVYDVVTNISVTFGVGIENDVTLDDTKSDFILLRDAKE  120
EssC5   LQANQAVNVENLAFYLCQDLHTQVYDVVTNISVTFGVGIENDVTLDDTKSDFILLRDAKE  120
EssC6   LQANQAVNVENLAFYLCQDLHTQVYDVVTNISVTFGVGIENDVTLDDTKSDFILLRDTKE  120
EssC7   LQANQAVNVENLAFYLCQDLHTQVYDVVTNISVTFGVGIENDVTLDDTKSDFILLRDTKE  120
EssC8   LQANQAVNVENLAFYLCQDLHTQVYDVVTNISVTFGVGIENDVTLDDTKSDFILLRDTKE  120

EssC1   KLFKLQVLNGEVYHNFSLVTEDCVLEPGDQLYTDGVTITIGKEDISVLAVKNRVTSKLAP  180
EssC2   KLFKLQVLNGEVYHNFSLVTEDCVLEPGDQLYTDGVTITIGKEDISVLAVKNRVTSKLAP  180
EssC3   GLFKLQVLNGEIYHNFSLVTEDCTVEPGDQLYTDGVTITIGKEDISVLAVKNRVTSKLAP  180
EssC4   KLFKLQVLNGEVYHNFSLVTEDCVLEPGDQLYTDGVTITIGKEDISVLAVKNRVTSKLAP  180
EssC5   KLFKLQVLNGEVYHNFSLVTEDCVLEPGDQLYTDGVTITIGKEDISVLAVKNRVTSKLAP  180
EssC6   GLFKLQVLNGEIYHNFSLVTEDCTVEPGDQLYTDGVTITIGKEDISVLAVKNRVTSKLAP  180
EssC7   GLFKLQVLNGEIYHNFSLVTEDCTVEPGDQLYTDGVTITIGKEDISVLAVKNRVTSKLAP  180
EssC8   GLFKLQVLNGEIYHNFSLVTEDCTVEPGDQLYTDGVTITIGKEDISVLAVKNRVTSKLAP  180

EssC1   LFAADNSFGEDYPDYHRSPRIIYRAPEEKISMAKPSSKPSKPTDGLIKIILPPLIMVAIT  240
EssC2   LFAADNSFGEDYPDYHRSPRIIYRAPEEKISMAKPSSKPSKPTDGLIKIILPPLIMVAIT  240
EssC3   LFAADNSFGEDYPDYHRSPRIIYRAPEEKISMAKPSSKPSKPTDGLIKIILPPLIMVAIT  240
EssC4   LFAADNSFGEDYPDYHRSPRIIYRAPEEKISMAKPSSKPSKPTDGLIKIILPPLIMVAIT  240
EssC5   LFAADNSFGEDYPDYHRSPRIIYRAPEEKISMAKPSSKPSKPTDGLIKIILPPLIMVAIT  240
EssC6   LFAADNSFGEDYPDYHRSPRIIYRAPEEKISMAKPSSKPSKPTDGLIKIILPPLIMVAIT  240
EssC7   LFAADNSFGEDYPDYHRSPRIIYRAPEEKISMAKPSSKPSKPTDGLIKIILPPLIMVAIT  240
EssC8   LFAADNSFGEDYPDYHRSPRIIYRAPEEKISMAKPSSKPSKPTDGLIKIILPPLIMVAIT  240

EssC1   VMISIFQPRGLYIIMTIAMSAVTITMAVLNYIKSRKKYKIDSKQRVESYDLYLKRKTKEL  300
EssC2   VMISIFQPRGLYIIMTIAMSAVTITMAVLNYIKSRKKYKIDSKQRVESYDLYLKRKTKEL  300
EssC3   VMISIFQPRGLYIIMTIAMSAVTITMAILNYIKSRKKYKIDSKQRVESYDLYLKRKTKEL  300
EssC4   VMISIFQPRGLYIIMTIAMSAVTITMAILNYIKSRKKYKIDSKQRVESYDLYLKRKTKEL  300
EssC5   VMISIFQPRGLYIIMTIAMSAVTITMAILNYIKSRKKYKIDSKQRVESYDLYLKRKTKEL  300
EssC6   VMISIFQPRGLYIIMTIAMSAVTITMAILNYIKSRKKYKIDSKQRVESYDLYLKRKTKEL  300
EssC7   VMISIFQPRGLYIIMTIAMSAVTITMAILNYIKSRKKYKIDSKQRVESYDLYLKRKTKEL  300
EssC8   VMISIFQPRGLYIIMTIAMSAVTITMAILNYIKSRKKYKIDSKQRVESYDLYLKRKTKEL  300

EssC1   HETSEKQRHALTYHYPDVTELEKMALRVDSRIYEKTMFHHDFLTFRVGRGDEASSFSVEF  360
EssC2   HETSEKQRHALTYHYPDVTELEKMALRVDSRIYEKTMFHHDFLTFRVGRGDETSSFSVEF  360
EssC3   HETSEKQRHALTYHYPDVTELEKMALRVDSRIYEKTMFHHDFLTFRVGRGDETSSFSVEF  360
EssC4   HETSEKQRHALTYHYPDVTELEKMALRVDSRIYEKTMFHHDFLTFRVGRGDEASSFSVEF  360
EssC5   HETSEKQRHALTYHYPDVTELEKMALRVDSRIYEKTMFHHDFLTFRVGRGDETSSFSVEF  360
EssC6   HETSEKQRHALTYHYPDVTELEKMALRVDSRIYEKTMFQHDFLTFRVGRGDEASSFSVEF  360
EssC7   HETSEKQRHALTYHYPDVTELEKMALRVDSRIYEKTMFHHDFLTFRVGRGDEASSFSVEF  360
EssC8   HETSEKQRHALTYHYPDVTELEKMALRVDSRIYEKTMFHHDFLTFRVGRGDEASSFSVEF  360

EssC1   QQEEFSQEKDELVEEAVKIKGQYLSINEVPVATDLMHGPVGYIGPRRLVLEQLQMLVMQT  420
EssC2   QQEEFSQEKDELVEEAVKIKGQYLSINEVPVATDLMHGPVGYIGPRRLVLEQLQMLVMQT  420
EssC3   QQEEFSQEKDELVEEAVKIKGQYLSINEVPVATDLMHGPVGYIGPRRLVLEQLQMLVMQT  420
EssC4   QQEEFSQEKDELVEEAVKIKGQYLSINEVPVATDLMHGPVGYIGPRRLVLEQLQMLVMQT  420
EssC5   QQEEFSQEKDELVEEAVKIKGQYLSINEVPVATDLMHGPVGYIGPRRLVLEQLQMLVMQT  420
EssC6   QQEEFSQEKDELVEEAVKIKGQYLSINEVPVATDLMHGPVGYIGPRRLVLEQLQMLVMQT  420
EssC7   QQEEFSQEKDELVEEAVKIKGQYLSINEVPVATDLMHGPVGYIGPRRLVLEQLQMLVMQT  420
EssC8   QQEEFSQEKDELVEEAVKIKGQYLSINEVPVATDLMHGPVGYIGPRRLVLEQLQMLVMQT  420
```

| | | |
|---|---|---|
| EssC1 | SLFHSYYDLQFITIFPEEEKADWDWMRWLPHANMRDVNVRGFVYHERSRDQVLNSLYQIL | 480 |
| EssC2 | SLFHSYYDLQFITIFPEEEKADWDWMRWLPHANMRDVNVRGFVYHERSRDQVLNSLYQIL | 480 |
| EssC3 | SLFHSYYDLQFITIFPEEEKADWDWMRWLPHANMRDVNVRGFVYHERSRDQVLNSLYQIL | 480 |
| EssC4 | SLFHSYYDLQFITIFPEEEKADWDWMRWLPHANMRDVNVRGFVYHERSRDQVLNSLYQIL | 480 |
| EssC5 | SLFHSYYDLQFITIFPEEEKVDWDWMRWLPHANMRDVNVRGFVYHERSRDQVLNSLYQIL | 480 |
| EssC6 | SLFHSYYDLQFITIFPEEEKADWDWMRWLPHANMRDVNVRGFVYHERSRDQVLNSLYQIL | 480 |
| EssC7 | SLFHSYYDLQFITIFPEEEKADWDWMRWLPHANMRDVNVRGFVYHERSRDQVLNSLYQIL | 480 |
| EssC8 | SLFHSYYDLQFITIFPEEEKPDWDWMRWLPHANMRDVNVRGFVYHERSRDQVLNSLYQIL | 480 |
| EssC1 | KERKQALTEQASKQEKLYFTPHYVVLITDEKLVLDHTVMEFFNEDPSELGVSLVFVQDVM | 540 |
| EssC2 | KERKQALTEQASKQEKLYFTPHYVVLITDEKLVLDHTVMEFFNEDPSELGVSLVFVQDVM | 540 |
| EssC3 | KERKQALTEQASKQEKLYFTPHYVVLITDEKLVLDHTVMEFFNEDPSELGVSLVFVQDVM | 540 |
| EssC4 | KERKQALTEQASKQEKLYFTPHYVVLITDEKLVLDHTVMEFFNEDPSELGVSLVFVQDVM | 540 |
| EssC5 | KERKQALTEQASKQEKLYFTPHYVVLITDEKLVLDHTVMEFFNEDPSELGVSLVFVQDVM | 540 |
| EssC6 | KERKQALTEQASKQEKLYFTPHYVVLITDEKLVLDHTVMEFFNEDPSELGVSLVFVQDVM | 540 |
| EssC7 | KERKQALTEQASKQEKLYFTPHYVVLITDEKLVLDHTVMEFFNEDPSELGVSLVFVQDVM | 540 |
| EssC8 | KERKQALTEQASKQEKLYFTPHYVVLITDEKLVLDHTVMEFFNEDPSELGVSLVFVQDVM | 540 |
| EssC1 | ESLPEHVKTVVDIRDAKSGNIILEQGDLVNRAFVPDHLPADFDKEVISRALAPLNHLQNL | 600 |
| EssC2 | ESLPEHVKTVVDIRDAKSGNIILEQGDLVNRAFVPDHLPADFDKEVISRALAPLNHLQNL | 600 |
| EssC3 | ESLPEHVKTVVDIRDAKSGNIILEQGDLVNRAFVPDHLPADFDKEVISRALAPLNHLQNL | 600 |
| EssC4 | ESLPEHVKTVVDIRDAKSGNIILEQGDLVNRAFVPDHLPADFDKEVISRALAPLNHLQNL | 600 |
| EssC5 | ESLPEHVKTVVDIRDAKSGNIILEQGDLVNRAFVPDHLPADFDKEVISRALAPLNHLQNL | 600 |
| EssC6 | ESLPEHVKTVVDIRDAKSGNIILEQGDLVNRAFVPDHLPADFDKEVISRALAPLNHLQNL | 600 |
| EssC7 | ESLPEHVKTVVDIRDAKSGNIILEQGDLVNRAFVPDHLPADFDKEVISRALAPLNHLQNL | 600 |
| EssC8 | ESLPEHVKTVVDIRDAKSGNIILEQGDLVNRAFVPDHLPADFDKEVISRALAPLNHLQNL | 600 |
| EssC1 | KNSIPESVTFLEMYGVERVEELNIAGRWAKNETYKSLAVPLGLRGKDDIVQLNLHEKAHG | 660 |
| EssC2 | KNSIPESVTFLEMYGVERVEELNIAGRWAKNETYKSLAVPLGLRGKDDIVQLNLHEKAHG | 660 |
| EssC3 | KNSIPESVTFLEMYGVERVEELNIAGRWAKNETYKSLAVPLGLRGKDDIVQLNLHEKAHG | 660 |
| EssC4 | KNSIPESVTFLEMYGVERVEELNIAGRWAKNETYKSLAVPLGLRGKDDIVQLNLHEKAHG | 660 |
| EssC5 | KNSIPESVTFLEMYGVERVEELNIAGRWAKNETYKSLAVPLGLRGKDDIVQLNLHEKAHG | 660 |
| EssC6 | KNSIPESVTFLEMYGVERVEELNIAGRWAKNETYKSLAVPLGLRGKDDIVQLNLHEKAHG | 660 |
| EssC7 | KNSIPESVTFLEMYGVERVEELNIAGRWAKNETYKSLAVPLGLRGKDDIVQLNLHEKAHG | 660 |
| EssC8 | KNSIPESVTFLEMYGVERVEELNIAGRWAKNETYKSLAVPLGLRGKDDIVQLNLHEKAHG | 660 |
| EssC1 | PHGLVAGTTGSGKSEIIQSYIISLGVNFHPYEVAFLLIDYKGGGMANLFKNMPHLLGTIT | 720 |
| EssC2 | PHGLVAGTTGSGKSEIIQSYIISLGVNFHPYEVAFLLIDYKGGGMANLFKNMPHLLGTIT | 720 |
| EssC3 | PHGLVAGTTGSGKSEIIQSYIISLGVNFHPYEVAFLLIDYKGGGMANLFKNMPHLLGTIT | 720 |
| EssC4 | PHGLVAGTTGSGKSEIIQSYIISLGVNFHPYEVAFLLIDYKGGGMANLFKNMPHLLGTIT | 720 |
| EssC5 | PHGLVAGTTGSGKSEIIQSYIISLGVNFHPYEVAFLLIDYKGGGMANLFKNMPHLLGTIT | 720 |
| EssC6 | PHGLVAGTTGSGKSEIIQSYIISLGVNFHPYEVAFLLIDYKGGGMANLFKNMPHLLGTIT | 720 |
| EssC7 | PHGLVAGTTGSGKSEIIQSYIISLGVNFHPYEVAFLLIDYKGGGMANLFKNMPHLLGTIT | 720 |
| EssC8 | PHGLVAGTTGSGKSEIIQSYIISLGVNFHPYEVAFLLIDYKGGGMANLFKNMPHLLGTIT | 720 |
| EssC1 | NLDGAQSMRALASIKAELQKRQRLFGEHDVNHINQYQKLYKQGKATEPMPHLFLISDEFA | 780 |
| EssC2 | NLDGAQSMRALASIKAELQKRQRLFGEHDVNHINQYQKLYKQGKATEPMPHLFLISDEFA | 780 |
| EssC3 | NLDGAQSMRALASIKAELQKRQRLFGEHDVNHINQYQKLYKQGKATEPMPHLFLISDEFA | 780 |
| EssC4 | NLDGAQSMRALASIKAELQKRQRLFGEHDVNHINQYQKLYKQGKATEPMPHLFLISDEFA | 780 |
| EssC5 | NLDGAQSMRALASIKAELQKRQRLFGEHDVNHINQYQKLYKQGKATEPMPHLFLISDEFA | 780 |
| EssC6 | NLDGAQSMRALASIKAELQKRQRLFGEHDVNHINQYQKLYKQGKATEPMPHLFLISDEFA | 780 |
| EssC7 | NLDGAQSMRALASIKAELQKRQRLFGEHDVNHINQYQKLYKQGKATEPMPHLFLISDEFA | 780 |
| EssC8 | NLDGAQSMRALASIKAELQKRQRLFGEHDVNHINQYQKLYKQGKATEPMPHLFLISDEFA | 780 |
| EssC1 | ELKSEQPEFMKELVSTARIGRSLGIHLILATQKPSGVVDDQIWSNSKFKLALKVQNASDS | 840 |
| EssC2 | ELKSEQPEFMKELVSTARIGRSLGIHLILATQKPSGVVDDQIWSNSKFKLALKVQNASDS | 840 |
| EssC3 | ELKSEQPEFMKELVSTARIGRSLGIHLILATQKPSGVVDDQIWSNSKFKLALKVQNASDS | 840 |
| EssC4 | ELKSEQPEFMKELVSTARIGRSLGIHLILATQKPSGVVDDQIWSNSKFKLALKVQNASDS | 840 |
| EssC5 | ELKSEQPEFMKELVSTARIGRSLGIHLILATQKPSGVVDDQIWSNSKFKLALKVQNASDS | 840 |
| EssC6 | ELKSEQPEFMKELVSTARIGRSLGIHLILATQKPSGVVDDQIWSNSKFKLALKVQNASDS | 840 |
| EssC7 | ELKSEQPEFMKELVSTARIGRSLGIHLILATQKPSGVVDDQIWSNSKFKLALKVQNASDS | 840 |
| EssC8 | ELKSEQPEFMKELVSTARIGRSLGIHLILATQKPSGVVDDQIWSNSKFKLALKVQNASDS | 840 |

```
EssC1    NEILKTPDAAEITLPGRSYLQVGNNEIYELFQSAWSGADYVPDKESTDYIDTTIYAINDL  900
EssC2    NEILKTPDAAEITLPGRSYLQVGNNEIYELFQSAWSGADYVPDKESTDYIDTTIYAINDL  900
EssC3    NEILKTPDAAEITLPGRSYLQVGNNEIYELFQSAWSGADYVPDKESTDYIDTTIYAINDL  900
EssC4    NEILKTPDAAEITLPGRSYLQVGNNEIYELFQSAWSGADYVPDKESTDYIDTTIYAINDL  900
EssC5    NEILKTPDAAEITLPGRSYLQVGNNEIYELFQSAWSGADYVPDKESTDYIDTTIYAINDL  900
EssC6    NEILKTPDAAEITLPGRSYLQVGNNEIYELFQSAWSGADYVPDKESTDYIDTTIYAINDL  900
EssC7    NEILKTPDAAEITLPGRSYLQVGNNEIYELFQSAWSGADYVPDKESTDYIDTTIYAINDL  900
EssC8    NEILKTPDAAEITLPGRSYLQVGNNEIYELFQSAWSGADYVPDKESTDYIDTTIYAINDL  900

EssC1    GQYDILTEDLSGLDKKDDLTKLPSELDAVIDHIHEYTEASGIEALPRPWLPPLEEQIFLP  960
EssC2    GQYDILTEDLSGLDKKDDLTKLPSELDAVIDHIHEYTEASGIEALPRPWLPPLPERIFAQ  960
EssC3    GQYDILTEDLSGLDKKDDLTKLPSELDAVIDHIHEYTEASGIEALPRPWLPPLEERISLE  960
EssC4    GQYDILTEDLSGLDKKDDLTKLPSELDAVIDHIHEYTEASGIEALPRPWLPPLPERIFAQ  960
EssC5    GQYDILTEDLSGLDKKDDLTKLPSELDAVIDHIHAYTEASGIEALPRPWLPPLPERIFAQ  960
EssC6    GQYDILTEDLSGLDKKDDLTKLPSELDAVIDHIHEYTEASGIEALPRPWLPPLPERIFAQ  960
EssC7    GQYDILTEDLSGLDKKDDLTKLPSELDAVIDHIHEYTEASGIEALPRPWLPPLPERIFAQ  960
EssC8    GQYDILTEDLSGLDKKDDLTKLPSELDAVIDHIHEYTEASGIEALPRPWLPPLEEQIFLP  960

EssC1    ELHQVITDELWSGEKQPLQATIGFLDIPQMQAQEPLTIDLAKDGHLAVFSSPGYGKSTFL  1020
EssC2    DLHPVNFEEAWKEPKKPLQATIGLLDQPELQAQVPLTLDLTKDGHVAVFSSPGFGKSTFL  1020
EssC3    SISTVDFEANWQKDEKDLELTLGVLDQPQLQAQNVLHWNLEKNGHMAVFSSPGFGKSTFM  1020
EssC4    DLHPVNFEEAWKEPKKPLQATIGLLDQPELQAQVPLTLDLTKDGHVAVFSSPGFGKSTFL  1020
EssC5    DLHPVNFEEAWKEPKKPLQATIGLLDQPELQAQVPLTLDLTKDGHVAVFSSPGFGKSTFL  1020
EssC6    DLHPVNFEEAWKEPKKPLQATIGLLDQPELQAQVPLTLDLTKDGHVAVFSSPGFGKSTFL  1020
EssC7    DLHPVNFEEAWKEPKKPLQATIGLLDQPELQAQVPLTLDLTKDGHVAVFSSPGFGKSTFL  1020
EssC8    ELHQVITDELWSGEKQPLQATIGFLDIPQMQAQEPLTIDLAKDGHLAVFSSPGYGKSTFL  1020

EssC1    QTITMDLARQHNPERLHIYLLDLGTNGLLPLKKLPHVADTIMVDEEIKIGKLIRRLTLEL  1080
EssC2    QSLVMDLARQHNPEQLHVYLLDFGTNGLLPLVDLPHIADTMMVDEVEKIQKFLRICLNEI  1080
EssC3    QTAIFDLARKNTPEFFHAYLLDFGTNGLLSLKGLPHVADTFSIDETEKTLKLVRLLSREI  1080
EssC4    QSLVMDLARQHNPEQLHVYLLDFGTNGLLPLIDLPHVADTMMVDEVEKVQKFVRRISEVM  1080
EssC5    QSLVMDLARQHNPEQLHVYLLDFGTNGLLPLIDLPHVADTIMIDEIEKARKFVKIVIREI  1080
EssC6    QSLVMDLARQHNPEQLHVYLLDFGTNGLLPLAKLPHTVDLLNIDNEEKLRKFTNRMNELI  1080
EssC7    QSLVMDLARQHNPEQLHVYLLDFGTNGLLSMKDVPHVADLMRLDEEEKITKLLKRVQNEI  1080
EssC8    QTITMDLARQHNPERLHIYLLDLGTNGLLPLKKLPHVADTIMVDEEIKIGKLIRRLYQEL  1080

EssC1    KERKQKLSKYGVASISMYEKASKEEVPAILLVIDAFDSVGEAP-YKDVFEKLIAQIAREG  1139
EssC2    KTRKKLLSQYRVANIEQYERASGKELPNIIIVLDNYDAVKDAG-LGDDFEKIITQITREG  1139
EssC3    KERKQLLSKFSVASLKMYEEISGDKKPIILLAIDNYDAIREVDEFVANLEPTIVQIAREG  1140
EssC4    KYRKKLLSKYRVANIEQYEKASKEEIPNIVIVLDNFDAVREAG-FGENFDKIMGQVSREG  1139
EssC5    KVRKKLLSEYRVANIEQYSQASGKNVANILVCLDNYDALREAG-FGDEFDKTMIQMAREG  1139
EssC6    SDRKKLLSKNAVANLSQFEEITKEVLPEVLVLIDNYDAIRETD-FSFGFESTLTQIAREG  1139
EssC7    ETRKKLLSEYSVASLEQYEKASQKQLPHILITLDGYDVVRDSD-LPPEFEKMLIQITREG  1139
EssC8    KIRKQKLSQYGVANISMYERASGEEIPSILLVIDAYDSISQAD-YKDSLEKIVAQIAREG  1139

EssC1    ASVGIHLVMSAVRQNAIRVQMLASIKHQIPLFMIEPGEARSIVGKTDLTIEELPGRGLVK  1199
EssC2    ASIGMFMIISASRHMSLRTQMATNIKQMIALYLIDKNDISSTVGRTDTPLEEYPGRALVK  1199
EssC3    ASLGIHLMITANNQNAMRLQLLSNIKTQIALHLNEKNEVSSIVGRSDYTIEELPGRGLIK  1200
EssC4    SSVGVFLATSASKYTSIKMQIVANIKLMVSLFIIDISDTRAIVGRTDLSVEELAGRGLVN  1199
EssC5    AALGIYLVTSASKQSSIRMQVMSSIKLQIALYLIDKSEVTSIVGRTDLILEELYGRGMVK  1199
EssC6    NSVGIHLVMSATRQNSMRQNLLANLKLQLALYMIEGNEVKSIVGQTKLTVAEFAGRGLVK  1199
EssC7    AAIGIHLALSAIRGAAMKPQMLMNFKLVVSLFNIDLSESRALIGRTDLTIEEIAGRGMVK  1199
EssC8    AGVGIHLVTSAVRQSAIRVQVLANIKNQISLYLIDQNEAKSIVGRTDLKIDEIPGRGLVK  1199

EssC1    LEEPTVFQTALPVCADGTLEIIEKIQAESEAMSSEWNGSRPAPIPMVPEVINMLEYLENK  1259
EssC2    LETVTTFQTTLPSQGEETIQQIEGIRNEAKLMREGWQGELPESVPMIPEKIKLADFCEWM  1259
EssC3    IEEPTLFQMALPNNGAEAIEIIKNNQDEAEKMTEIWTGQKPQCIPMVPETLGFTHFTDHM  1260
EssC4    QDGVSVMQSILPADGEDVLEQISNLQSEAKSMRCAWKGHLPEDIPMMPDELTFTNFITKE  1259
EssC5    VGSQAIFQTTLPTKGREIVEQINNLTEEIAVMKESWNGECPDSIPVMPEVLSISDFVGRK  1259
EssC6    LEDPTLFQASFPTRSNSVTEHLECLDKEILQMSKHGIYKVVPKIPMLPDKIGLDELISKK  1259
EssC7    LEHPTIFQVNTPTEGDDILETIANIKIEAKKMEESWNGERPEAIPVVPEKMEYNAFATMP  1259
EssC8    LEDPTSFQAALPVAGSDTLEIIDKVQKECQVMRDLWKGILPISIPMVPEEFSIKQFLSMD  1259
```

```
EssC1    QVQKSLSE-GKTPIAVDFEDVLPVNLDVQTDGNTLVLTDNADILERTM----VSLTEL-I 1313
EssC2    ETKEVVAN-GNIPIGVDFEYVKPVGIDYKQTGVLIIADATGRNLVSIN----KNIANS-L 1313
EssC3    ETKKMLELKSILPIGLEYEYASPVGVSLEQHNLAIIMPTVELVQQII-----TNSSHL-L 1314
EssC4    SVK-NINN-NEIAVGLDFEVVEGVRWDLQKLDNILILGGSQAGKTTMQHTLLKGIERANL 1317
EssC5    TTKDNIQS-EILPLGLEYENIEAVGLDFKLKQNLLIVSDSNIKLNVVQ----QSIVKSGQ 1314
EssC6    SVQQSLAN-HLFVIGLDEEAVESFAVNKINQLPLLVTGENGFDISNAI----KQFIDLES 1314
EssC7    YVQKLMES-NLVPIGFDMETALPVGLDLNKQRIWLITGSETDLNENTFHAMIKGIQKI-- 1316
EssC8    KTKEYLDVYGELPIGIDFEYAEAVRVALYKQNLVVVTTTSQELIEKL-----NSLINI-L 1313

EssC1    GQNMEVDIALYDNSSNRFQQYRNN--VNIYAGEETAMNSASAQMVSVLEAREDGWKEIQR 1371
EssC2    VQNPNISLLISDTANFDLVEMSTK--ANLYSNNTEDLKQNMKKVLSIYQSRTEKADELRK 1371
EssC3    VKKELRELVVFDTPSMELMKLKEL--PLTYITNSERIEGKIDILYGAFKTAEQAYLEELQ 1372
EssC4    MKRDTNIIHIFDTNRGKLKEYKEKEIVANYIVDKDAYGEFFNGLSELLEYRFNELLRIRD 1377
EssC5    SVKSSYQLLILDKISQPLKDVSVW--ANGYVSKLEEYEKFTSMMEEEFATRKD------- 1365
EssC6    KDFESKEIIVIDNMKYDFSKYQND--IYIYGNNKTFMETIIQNSMNEYEERRKVVTRLRE 1372
EssC7    ----GQKLHLVDLSNYNLANLNSE--NTNFISKPETFQEVIKEVYAENNVREEAVRKCSP 1370
EssC8    TKKSDVETTVIDQKAMELISINDN--VSDYITDIESVETFIEDVYDDYREREAAFQEEIQ 1371

EssC1    ASNGEITLKMYVEELRPMYVLLTDSIYAAEQMSLEARKNIVRLIENGPKLGIYFVTGSQT 1431
EssC2    -AGDNLGVTAMIQGLKDIVVFVGDYDILLKALDEEDQKTLIDMINNGKTYGCYLILSGSI 1430
EssC3    -SNNTITKEEYFKNVPPIVVVTYSTVQLYNQLSSNAQKQLLEMLKSDGKMGVEFIIGNDL 1431
EssC4    -MDGEKAGEEFIQGEQKHFIFIQNIGDFGNGLPTEVVKKLVYILDNNSKLGIHFIISGTS 1436
EssC5    -----LDSKALLKDTEGWIVLISDLAEFIQH-SSITIERMKEFVENGPNLNIFFIISGMQ 1419
EssC6    ENRTRIEIEEQLANMPEILLFITDLSSVIESVEAKYLDMLEKMLGDAAVYKVFFTIGASL 1432
EssC7    -----PELNAYYQTIETINIAV-DTLEVFSKVDSETQRMIQEMMTSRIQTNINIIIQVDI 1424
EssC8    QSNGQTSPIDFYKKYKAKCIIFNGFIQIQNDLSTRTQGQLLELLKQEGKTGIWFVIGNEL 1431

EssC1    GILYRARDEISGELR-KQKTGILLGRISDQNILSLINNIYKEALLAPYEAYYIKQGQYEK 1490
EssC2    QAF-KGYGDLNKAVQ-AQRNYIFFDRLSNSSPFSFSFNIYREQELLPEEGYVIAGQEWQK 1488
EssC3    GSMAKEYSPIGDAIR-NSKQVLLGARFADQAIYSPTIRIVQEKPLAPQELYLLLEGNAEK 1490
EssC4    NNFGQNYSDFTNRVK-QINSGIIIAGYNEQSIVRMDNVNMYSPKLGAGDAYFVDNGRATR 1495
EssC5    TSIEYGLNPIEKFVKTTIRTGLVAMKNADQGIFK-GNYISNEQPLEKFETYFYVDSTRQK 1478
EssC6    GSLS-GYDSLSKTVK-AMKNVLLFTDVKQQSVINVVSKSGILKPLKPFEAYYIDNKIYSK 1490
EssC7    SDIGRDGSSVGRELR-NVTNLILSMKIKNQTTFAQDIRSYDEPDLPPRTSYYINGKLASK 1483
EssC8    GIIGKAYGDLANKVK-ESKLVILAMRVTDQTVFSTINRIVQEQQLRKGEVYLIANNQPQK 1490

EssC1    IKLIAPNQ-- 1498
EssC2    IKIAKYEEEV 1498
EssC3    LKIPKG---- 1496
EssC4    IRMPKH---- 1501
EssC5    IKLPN----- 1483
EssC6    VKVAYVKEEE 1500
EssC7    FRVLSKS--- 1490
EssC8    VKIGV----- 1495
```

| | EssC1 | EssC2 | EssC3 | EssC4 | EssC5 | EssC6 | EssC7 | EssC8 |
|---|---|---|---|---|---|---|---|---|
| EssC1 | 100.00 | | | | | | | |
| EssC2 | 77.87 | 100.00 | | | | | | |
| EssC3 | 77.24 | 76.76 | 100.00 | | | | | |
| EssC4 | 77.44 | 80.64 | 75.55 | 100.00 | | | | |
| EssC5 | 78.12 | 81.49 | 77.50 | 81.16 | 100.00 | | | |
| EssC6 | 76.89 | 77.56 | 76.62 | 77.64 | 78.06 | 100.00 | | |
| EssC7 | 77.52 | 79.73 | 77.22 | 79.03 | 79.86 | 78.72 | 100.00 | |
| EssC8 | 82.80 | 77.88 | 79.72 | 77.61 | 78.18 | 77.36 | 78.22 | 100.00 |

Figure S2. Sequence alignment of the eight EssC variants found in *L. monocytogenes*. The shaded lines above the sequence indicate the approximate domain boundaries of EssC; red – FHA domains; grey – transmembrane domains; yellow – D0; blue – D1; orange – D2; green D3. The boxes indicate the percentage identity between each variant.

A

```
Toxin_D          MSRIDIGEIQAFLYQLRAANEPGRKTIQSIKAAVTKYVGDNSLKGKAVDASKNYYQMTYF 60
Toxin_E          MSRIDIGEIQAFLYQLRAANEPGRKTIQSIKAAVTKYVGDNSLKGKAVDASKNYYQMTYF 60
Toxin_F          MSRIDIGEIQAFLYQLRAANEPGRKTIQSIKAAVTKYVGDNSLKGKAVDASKNYYQMTYF 60
Toxin_G          MSRIDIGEIQAFLYQLRAANESGRKTIQSIKTAVTKYVGDNSLKGKAVDTSKNYYQMTYF 60
Toxin_Omicron    MSRIDIGEIQVFLYQLRAANESGRKTIQSIKTAVTKYVGDNSLKGKAVDASKTYYQMTYF 60

Toxin_D          PLCDAIIEAMDESEERLGQYIQDFHAEVDSSPDAKIDADGLYELGKMIDRIESKKEALAQ 120
Toxin_E          PLCDAIIEAMDESEERLGQYIQDFHAEVDSSPDAKIDADGLYELGKMIDRIESKKEALAQ 120
Toxin_F          PLCDAIIEAMDESEERLGQYIQDFHAEVDSSPDAKIDADGLYELGKMIDRIESKKEALAQ 120
Toxin_G          PLCDAIIEAMDESEERLGQYIQDFHAEVDSSPDAKIDADGLYELGKMIDRIESKKEALAQ 120
Toxin_Omicron    PLCDAIIEAMDESEERLGQYIQDFHAEVDSSPDAKIDADGLYELGKMIDRIESKKETLAQ 120

Toxin_D          RMNSGTEGQMQNYRSQLAIAYKQENILEKYLSFEQSHANFFDHLIDLVQAVQQTIRELQS 180
Toxin_E          RMNSGTEGQMQNYRSQLAIAYKQENILEKYLSFEQSHANFFDHLIDLVQAVQQTIRELQS 180
Toxin_F          RMNSGTEGQMQNYRSQLAIAYKQENILEKYLSFEQSHASFFDHLIDLVQAVQQTIRELQS 180
Toxin_G          RMNSGTEGQMQNYRSQLAIAYKQENILEKYLSFEQSHASFFDHLIDLVQAVQQTIRELQS 180
Toxin_Omicron    RMNSGTEGQMQNYRSQLAIAYKQENILEKYLSFEQSHASFFDHLIDLVQAVQQTIRELQS 180

Toxin_D          NIQFNSQTGTYDLSKLNSATVSRMQQALNKSRGIKEDIIKELRDYTVLAVVYLDSNGKEQ 240
Toxin_E          NIQFNSQTGTYDLSKLNSATVSRMQQALNKSRGIKEDIIKELRDYTVLAVVYLDSNGKEQ 240
Toxin_F          NIQFNSQTGTYDLSKLNHATVSHMQQALNKARGIKEDIIKELQDYTVLAVVYLDSNGKEQ 240
Toxin_G          NIQFNSQTGTYDLSKLNSATVSRMQQALNKSRGIKEDIIKELRDYTVLAVVYLDSNGKEQ 240
Toxin_Omicron    NIQFNSQTGTYDLSKLNSATVSRMQQALNKSRGIKEDIIKELRDYTVLAVVYLDSNGKEQ 240

Toxin_D          VMWLLERDGVGVENAELKAYLTENGKYLNPEDYTIITNEELNKKINQSWRDGVYYLNGNK 300
Toxin_E          VMWLLERDGLGVENAELKAYLEKNGKYLNPEDYSIITNEDLNKKINKAWRDGVYYLNGNK 300
Toxin_F          VMWLLERDGLGVENAELKAYLEKNGKYLNPEDYSIITNEDLNKKINKAWRDGVYYLNGNK 300
Toxin_G          VMWLLERDGVGVENAELKAYLTENGKYLNPEDYTIITNEELNKKINQSWRDGVYYLNGNK 300
Toxin_Omicron    VMWLLERDGVGVENAELKAYLTENGKYLNPEDYTIITNEELNKKINQSWRDGVYYLNGNK 300

Toxin_D          YDGLTGGVLSTSAYVEAGKGYIDKSGFADVVLGLGLSTAAIRGSLVYKKVN--------- 351
Toxin_E          YDGLTGGVLSTSAYVEAGKGYIDKSGLADVVLGLGLSTAAIRNATVIKSKS--------- 351
Toxin_F          YDGLTGGILSTSAYVEAGKGFIDKSGLADVVLGLGLSTAAIRGSVTFGKKNKLKGYDYLD 360
Toxin_G          YDGLTGGVLSTSAYVEAGKGYIDKSGLADVVLGLGLSTAAIRGSNTFGKKQS------VS 354
Toxin_Omicron    YDGLTGGVLSTSAYVEAGKGYIDKSGLADVVLGLGLSTAAIRGSMTFGKKHKLKGYDYLD 360

Toxin_D          ---GPYKQNIK-NRYPNEVQQGKIF--DYTLENGQVKIRDGIKEVDFIIDLQGNLKVGRG 405
Toxin_E          ------------SQLDYVTNTGRL-------DGTLYSSKDLVLLENYLKKRGIELKIGDK 392
Toxin_F          DLLGDLNNKVKIKKYDSAESVNKYWH-QQNYDQPPYTPKTPVQDLELLIETKFVRVYDGG 419
Toxin_G          NVPNSSKNSAQYQKYKNELMKGDIL--E---NSKPIILGSDLKDSKVVSALTKNGSSMSD 409
Toxin_Omicron    DQLGSMKSHVKVNKYESSENANNWWKVEKGYDNPPYTSKTVVQDIRLLSDETFVRVYDGD 420

Toxin_D          HAHLSNGGDVQAA-GKLKVDSNGNVRRITNESGHYTP--TLGQAKNYQ-----QIFENTG 457
Toxin_E          YLPANKGGAFNAETGTIILKDNPTRYEVLHEVSHYIQYKNIGKGAYKNLPRAGKTFDTKQ 452
Toxin_F          SSKLHGGWLMKAEDIKGLTPTQIKDKFALPNMPKFVGEVTLPKGSNIRMGEVNPLFENKG 479
Toxin_G          WAKMESTYSYETSMGRGKIHFYKNLK--TGEINFYDVKMKVPIPKDLK-------IRNQ- 459
Toxin_Omicron    VSGLKGGWLMRAEDIRGLTPKQIQAKFALPAEPIYIGEVSLPKGSKLRIGEVAKNFGHKG 480

Toxin_D          ----INTKNAWLETYQLDVTKSG--YVDLAKL----KRIDSVKLK 492
Toxin_E          ----FNAPEQFVYDMLSNNTRRWKSFTEAERMHANWYINNFGGIR 493
Toxin_F          GGIQFDLKGQFIGE-----------FKELG------KISEWGGLK 507
Toxin_G          -----LTDDFWIVDLDNN-------FIPKG-------------VR 479
Toxin_Omicron    GGIQFDLKGQYIGD-----------YKEVG------KIIEWGK-- 506
```

B

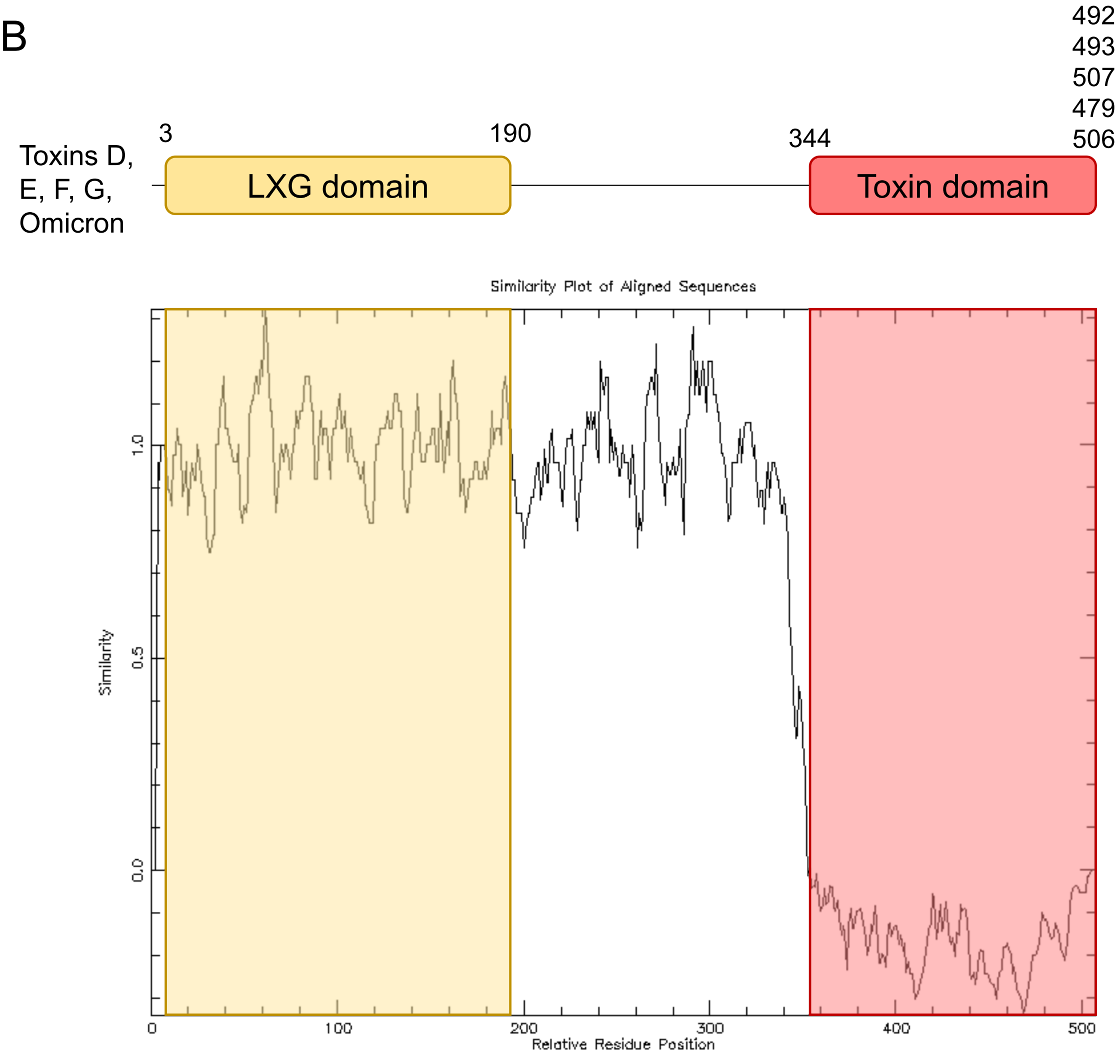

Figure S3. LXG toxins D, E, F, G and Omicron share a common N-terminal sequence. A. Sequence alignment of the five toxins. B. Analysis of the same five toxins using plotcon. The predicted boundary of the LXG domain is indicated.

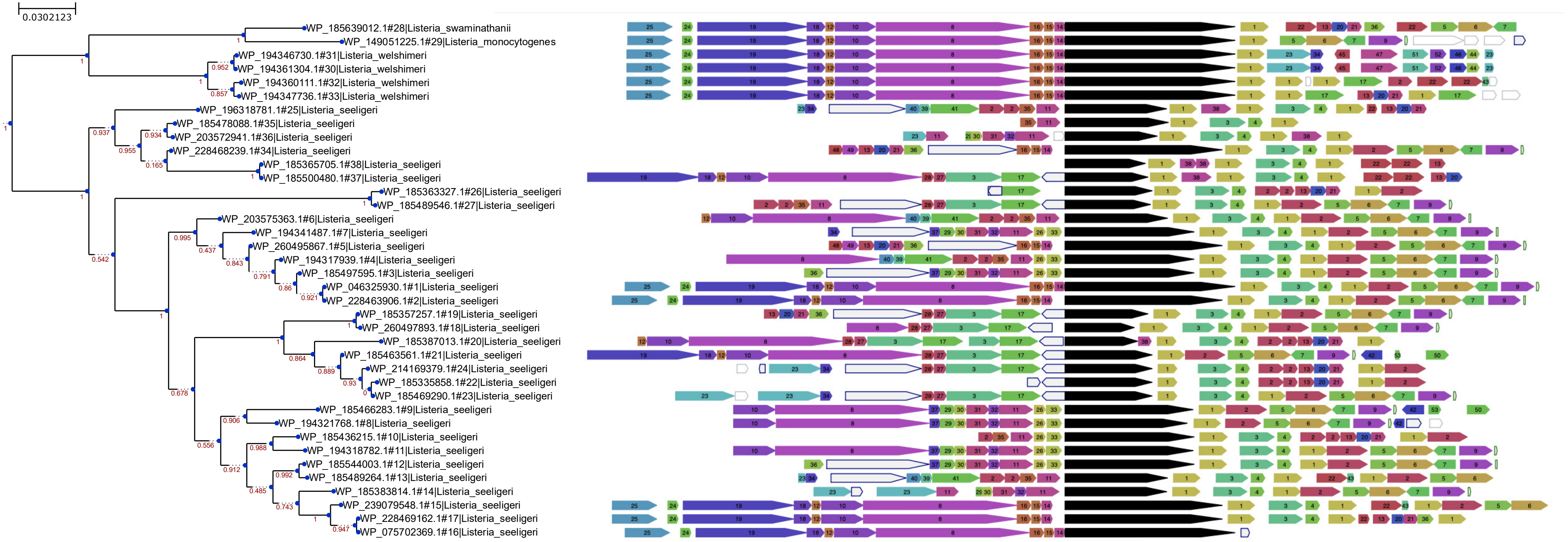

Figure S4. Genomic organisation of *lrhA* genes in non-monocytogenes species of *Listeria*. WebFlags was used to generate a genetic neighbourhood analysis of all LrhA homologues found in non-monocytogenes species. The tree was generated by webFlaGs using the ETE3 toolkit.

1(1) WP_203577659.1 hypothetical protein
1(1) WP_228463713.1 hypothetical protein
1(1) WP_185436222.1 hypothetical protein
1(1) WP_196318785.1 hypothetical protein
1(1) WP_185489543.1 hypothetical protein
1(1) WP_185469288.1 hypothetical protein
1(1) WP_194341489.1 hypothetical protein
1(1) WP_185415325.1 hypothetical protein
1(2) WP_185363328.1 hypothetical protein
1(2) WP_185365696.1 hypothetical protein
1(3) WP_185383815.1 hypothetical protein
1(2) WP_221633580.1 hypothetical protein
1(2) WP_185357268.1 hypothetical protein
1(1) WP_196318784.1 hypothetical protein
1(2) WP_185544005.1 hypothetical protein
1(1) WP_185387006.1 hypothetical protein
1(6) WP_046325931.1 hypothetical protein
1(1) WP_221638402.1 hypothetical protein
1(1) WP_185310098.1 hypothetical protein
1(1) WP_203572943.1 hypothetical protein
1(1) WP_194347735.1 hypothetical protein
1(1) WP_185383817.1 hypothetical protein
1(2) WP_194358207.1 hypothetical protein
1(2) WP_185365702.1 hypothetical protein
1(2) WP_194346731.1 hypothetical protein
1(1) WP_185466281.1 hypothetical protein
1(3) WP_185335851.1 hypothetical protein
1(1) WP_194341491.1 hypothetical protein
1(1) WP_185489260.1 hypothetical protein
1(1) WP_185489262.1 hypothetical protein
1(1) WP_185478090.1 hypothetical protein
1(1) WP_196318782.1 hypothetical protein
1(1) WP_185639011.1 hypothetical protein
1(1) WP_239077241.1 hypothetical protein
1(1) WP_194347732.1 hypothetical protein
1(1) WP_194347734.1 hypothetical protein
1(3) WP_185415324.1 hypothetical protein
1(1) WP_228469160.1 hypothetical protein
1(1) WP_185436216.1 hypothetical protein
1(2) WP_185365691.1 hypothetical protein
1(1) WP_203577657.1 hypothetical protein
1(1) WP_203575366.1 hypothetical protein
1(4) WP_228463346.1 hypothetical protein
1(1) WP_194360113.1 hypothetical protein
1(1) WP_203575364.1 hypothetical protein
1(4) WP_046325932.1 hypothetical protein
1(1) WP_149051226.1 hypothetical protein
1(1) WP_194358203.1 hypothetical protein
1(1) WP_194360112.1 hypothetical protein
1(1) WP_194317940.1 hypothetical protein

2(1) WP_221637190.1 hypothetical protein
2(2) WP_250210513.1 hypothetical protein
2(2) WP_221637815.1 hypothetical protein
2(1) WP_243285277.1 hypothetical protein
2(1) WP_214169381.1 hypothetical protein
2(1) WP_185489266.1 hypothetical protein
2(1) WP_228472257.1 hypothetical protein
2(1) WP_260448395.1 hypothetical protein
2(1) WP_185455567.1 hypothetical protein
2(1) WP_228474727.1 hypothetical protein
2(1) WP_221639762.1 hypothetical protein
2(4) WP_003745060.1 hypothetical protein
2(3) WP_012984615.1 hypothetical protein
2(1) WP_228463232.1 hypothetical protein
2(3) WP_012984616.1 hypothetical protein
2(3) WP_221633573.1 hypothetical protein
2(1) WP_185387010.1 hypothetical protein
2(1) WP_221645729.1 hypothetical protein
2(1) WP_185436218.1 hypothetical protein
2(1) WP_185455569.1 hypothetical protein
2(1) WP_221770661.1 hypothetical protein
2(1) WP_221642425.1 hypothetical protein
2(1) WP_260444442.1 hypothetical protein
2(1) WP_185363330.1 hypothetical protein
2(2) WP_221770127.1 hypothetical protein
2(1) WP_221645600.1 hypothetical protein
2(1) WP_221638398.1 hypothetical protein
2(1) WP_185335855.1 hypothetical protein
2(1) WP_221635928.1 hypothetical protein
2(1) WP_221639041.1 hypothetical protein
2(1) WP_228463708.1 hypothetical protein
2(1) WP_239077615.1 hypothetical protein
2(1) WP_038408420.1 hypothetical protein
2(1) WP_228478965.1 hypothetical protein

3(1) WP_185340229.1 hypothetical protein
3(2) WP_185463563.1 T7SS effector LXG polymorphic toxin
3(1) WP_185436217.1 hypothetical protein
3(1) WP_239077240.1 hypothetical protein
3(1) WP_228478964.1 hypothetical protein
3(1) WP_203575365.1 hypothetical protein
3(1) WP_260448413.1 hypothetical protein
3(1) WP_260442961.1 hypothetical protein
3(1) WP_185387014.1 T7SS effector LXG polymorphic toxin
3(1) WP_185478089.1 hypothetical protein
3(1) WP_185365694.1 hypothetical protein
3(1) WP_185489263.1 hypothetical protein
3(1) WP_250210512.1 hypothetical protein
3(1) WP_239079539.1 hypothetical protein
3(1) WP_228463347.1 hypothetical protein
3(1) WP_228463449.1 hypothetical protein
3(4) WP_185357254.1 T7SS effector LXG polymorphic toxin
3(1) WP_250209931.1 hypothetical protein
3(1) WP_260444553.1 hypothetical protein
3(1) WP_260443318.1 hypothetical protein
3(2) WP_228469720.1 hypothetical protein
3(1) WP_003750480.1 hypothetical protein
3(1) WP_228465423.1 hypothetical protein
3(1) WP_260448424.1 hypothetical protein
3(1) WP_260447974.1 hypothetical protein
3(6) WP_228463507.1 hypothetical protein
3(1) WP_214169378.1 T7SS effector LXG polymorphic toxin

4(3) WP_185357265.1 hypothetical protein
4(12) WP_003745059.1 hypothetical protein
4(4) WP_003750481.1 hypothetical protein
4(5) WP_185335856.1 hypothetical protein
4(1) WP_185451814.1 hypothetical protein
4(1) WP_185365693.1 hypothetical protein
4(1) WP_203572942.1 hypothetical protein
4(1) WP_185489544.1 hypothetical protein

5(2) WP_185544006.1 isocitrate lyase/phosphoenolpyruvate mutase family protein
5(1) WP_203575398.1 isocitrate lyase/phosphoenolpyruvate mutase family protein
5(1) WP_185463375.1 isocitrate lyase/phosphoenolpyruvate mutase family protein
5(1) WP_185489284.1 isocitrate lyase/phosphoenolpyruvate mutase family protein
5(1) WP_185639005.1 isocitrate lyase/phosphoenolpyruvate mutase family protein
5(1) WP_185469333.1 isocitrate lyase/phosphoenolpyruvate mutase family protein
5(1) WP_185489551.1 isocitrate lyase/phosphoenolpyruvate mutase family protein
5(4) WP_046325933.1 isocitrate lyase/phosphoenolpyruvate mutase family protein
5(1) WP_203577660.1 isocitrate lyase/phosphoenolpyruvate mutase family protein
5(1) WP_149051227.1 isocitrate lyase/phosphoenolpyruvate mutase family protein
5(1) WP_194341498.1 isocitrate lyase/phosphoenolpyruvate mutase family protein
5(1) WP_185383818.1 isocitrate lyase/phosphoenolpyruvate mutase family protein
5(1) WP_139591395.1 isocitrate lyase/phosphoenolpyruvate mutase family protein
5(1) WP_185463620.1 isocitrate lyase/phosphoenolpyruvate mutase family protein
5(1) WP_185415376.1 isocitrate lyase/phosphoenolpyruvate mutase family protein
5(3) WP_185335869.1 isocitrate lyase/phosphoenolpyruvate mutase family protein

6(1) WP_203577661.1 trifunctional transcriptional activator/DNA repair protein Ada/methylated-DNA--
6(7) WP_003745063.1 trifunctional transcriptional activator/DNA repair protein Ada/methylated-DNA--
6(1) WP_194341492.1 trifunctional transcriptional activator/DNA repair protein Ada/methylated-DNA--
6(1) WP_185461316.1 trifunctional transcriptional activator/DNA repair protein Ada/methylated-DNA--
6(1) WP_185363332.1 trifunctional transcriptional activator/DNA repair protein Ada/methylated-DNA--
6(1) WP_185463559.1 trifunctional transcriptional activator/DNA repair protein Ada/methylated-DNA--

6(1) WP_221771473.1 trifunctional transcriptional activator/DNA repair protein Ada/methylated-DNA--
6(3) WP_185357272.1 trifunctional transcriptional activator/DNA repair protein Ada/methylated-DNA--
6(1) WP_185335850.1 trifunctional transcriptional activator/DNA repair protein Ada/methylated-DNA--
6(1) WP_149051228.1 trifunctional transcriptional activator/DNA repair protein Ada/methylated-DNA--
6(2) WP_185383819.1 trifunctional transcriptional activator/DNA repair protein Ada/methylated-DNA--
6(2) WP_185469976.1 trifunctional transcriptional activator/DNA repair protein Ada/methylated-DNA--

7(3) WP_185357275.1 pentapeptide repeat-containing protein
7(2) WP_185455618.1 pentapeptide repeat-containing protein
7(4) WP_046325947.1 pentapeptide repeat-containing protein
7(1) WP_194332042.1 pentapeptide repeat-containing protein
7(1) WP_185469332.1 pentapeptide repeat-containing protein
7(1) WP_185383872.1 pentapeptide repeat-containing protein
7(3) WP_185415377.1 pentapeptide repeat-containing protein
7(2) WP_185469980.1 pentapeptide repeat-containing protein
7(1) WP_185350592.1 pentapeptide repeat-containing protein
7(1) WP_149051229.1 pentapeptide repeat-containing protein
7(1) WP_185639003.1 pentapeptide repeat-containing protein

8(1) WP_194347737.1 type VII secretion protein EssC
8(1) WP_194360110.1 type VII secretion protein EssC
8(1) WP_194318783.1 type VII secretion protein EssC
8(2) WP_046325927.1 type VII secretion protein EssC
8(1) WP_185466285.1 type VII secretion protein EssC
8(1) WP_185436428.1 type VII secretion protein EssC
8(1) WP_185387015.1 type VII secretion protein EssC
8(1) WP_203575361.1 type VII secretion protein EssC
8(1) WP_194321762.1 type VII secretion protein EssC
8(1) WP_260497892.1 FtsK/SpoIIIE domain-containing protein, partial
8(1) WP_185639016.1 type VII secretion protein EssC
8(1) WP_203577655.1 type VII secretion protein EssC
8(1) WP_149051221.1 type VII secretion protein EssC
8(1) WP_185364860.1 type VII secretion protein EssC
8(2) WP_075702371.1 type VII secretion protein EssC
8(1) WP_194346729.1 type VII secretion protein EssC
8(1) WP_185463564.1 type VII secretion protein EssC
8(1) WP_194361305.1 type VII secretion protein EssC

9(1) WP_149051269.1 2-hydroxyacid dehydrogenase family protein
9(1) WP_185387003.1 2-hydroxyacid dehydrogenase family protein
9(3) WP_185357278.1 2-hydroxyacid dehydrogenase family protein
9(1) WP_185455580.1 hydroxyacid dehydrogenase
9(2) WP_185415326.1 2-hydroxyacid dehydrogenase family protein
9(1) WP_185463558.1 hydroxyacid dehydrogenase
9(1) WP_139590304.1 2-hydroxyacid dehydrogenase family protein
9(3) WP_046325934.1 2-hydroxyacid dehydrogenase family protein
9(1) WP_185383820.1 hydroxyacid dehydrogenase
9(1) WP_185469287.1 hydroxyacid dehydrogenase
9(1) WP_075703054.1 hydroxyacid dehydrogenase
9(2) WP_185469977.1 hydroxyacid dehydrogenase
9(1) 2-hydroxyacid dehydrogenase family protein

10(1) WP_194321760.1 type VII secretion protein EssB
10(2) WP_075702372.1 type VII secretion protein EssB
10(1) WP_185433816.1 type VII secretion protein EssB
10(1) WP_139590297.1 type VII secretion protein EssB
10(1) WP_194318784.1 type VII secretion protein EssB
10(1) WP_012582211.1 type VII secretion protein EssB
10(1) WP_185307483.1 type VII secretion protein EssB
10(1) WP_185463565.1 type VII secretion protein EssB
10(1) WP_203575360.1 type VII secretion protein EssB
10(1) WP_185639017.1: type VII secretion protein EssB
10(1) WP_194360109.1 type VII secretion protein EssB
10(1) WP_011700924.1 type VII secretion protein EssB
10(2) WP_046325926.1 type VII secretion protein EssB
10(1) WP_185363042.1 type VII secretion protein EssB
10(2) WP_185337267.1 type VII secretion protein EssB

11(3) WP_185478087.1 hypothetical protein
11(3) WP_185365747.1 hypothetical protein
11(2) WP_185383809.1 hypothetical protein
11(1) WP_185383813.1 hypothetical protein
11(1) WP_194341483.1 hypothetical protein
11(1) WP_203572939.1 hypothetical protein
11(1) WP_185489265.1 hypothetical protein
11(2) WP_185436213.1 hypothetical protein
11(1) WP_185455574.1 hypothetical protein
11(2) WP_185497592.1 hypothetical protein

12(2) WP_003759563.1 EsaB/YukD family protein
12(1) WP_194347738.1 EsaB/YukD family protein
12(9) WP_003745045.1 EsaB/YukD family protein
12(1) WP_185442773.1 EsaB/YukD family protein
12(2) WP_003728973.1 EsaB/YukD family protein

13(1) WP_185363331.1 hypothetical protein
13(1) WP_185630933.1 hypothetical protein
13(1) WP_185337274.1 hypothetical protein
13(2) WP_185335854.1 hypothetical protein
13(2) WP_185415320.1 hypothetical protein
13(1) WP_196318786.1 hypothetical protein
13(2) WP_185365686.1 hypothetical protein
13(1) WP_185436219.1 hypothetical protein
13(1) WP_185387009.1 hypothetical protein
13(1) WP_185639009.1 hypothetical protein
13(1) WP_194358205.1 hypothetical protein

14(1) WP_185639013.1 hypothetical protein
14(7) WP_046325929.1 hypothetical protein
14(1) WP_149051224.1 hypothetical protein
14(4) WP_185331649.1 hypothetical protein

15(1) WP_185325795.1 hypothetical protein
15(5) WP_075702370.1 hypothetical protein
15(1) WP_185639014.1 hypothetical protein
15(2) WP_003745050.1 hypothetical protein
15(3) WP_185331648.1 hypothetical protein
15(1) WP_149051223.1 hypothetical protein

16(1) WP_149051222.1 hypothetical protein
16(4) WP_185325794.1 hypothetical protein
16(1) WP_185639015.1 hypothetical protein
16(7) WP_046325928.1 hypothetical protein

17(2) WP_185463562.1 hypothetical protein
17(1) WP_185335969.1 hypothetical protein
17(1) WP_194347731.1 hypothetical protein
17(1) WP_194347733.1 hypothetical protein
17(1) WP_194360114.1 hypothetical protein
17(7) WP_185340230.1 hypothetical protein

18(1) WP_185463566.1 type VII secretion protein EssA
18(1) WP_185639018.1 type VII secretion protein EssA
18(1) WP_194361306.1 type VII secretion protein EssA
18(2) WP_046325925.1 type VII secretion protein EssA
18(1) WP_070214792.1 type VII secretion protein EssA
18(3) WP_003750475.1 type VII secretion protein EssA
18(3) WP_011700923.1 type VII secretion protein EssA
18(1) WP_185337266.1 type VII secretion protein EssA

19(2) WP_046325924.1 type VII secretion protein EsaA
19(1) WP_203577654.1 type VII secretion protein EsaA
19(1) WP_185639019.1 type VII secretion protein EsaA
19(1) WP_185463567.1 type VII secretion protein EsaA
19(1) WP_194360108.1 type VII secretion protein EsaA
19(2) WP_075702373.1 type VII secretion protein EsaA
19(1) WP_185309473.1 type VII secretion protein EsaA
19(1) WP_149051220.1 type VII secretion protein EsaA
19(1) WP_185364862.1 type VII secretion protein EsaA
19(1) WP_185327606.1 type VII secretion protein EsaA
19(1) WP_194346728.1 type VII secretion protein EsaA

20(1) WP_185304653.1 hypothetical protein
20(1) WP_185387008.1 hypothetical protein
20(1) WP_194358204.1 hypothetical protein
20(2) WP_185415321.1 hypothetical protein
20(1) WP_185436220.1 hypothetical protein
20(1) WP_185468865.1 hypothetical protein
20(1) WP_185365684.1 hypothetical protein
20(1) WP_185639008.1 hypothetical protein
20(2) WP_185335853.1 hypothetical protein
20(1) WP_185337275.1 hypothetical protein
20(1) WP_214169382.1 hypothetical protein

21(3) WP_185337276.1 DUF4064 domain-containing protein
21(2) WP_185335852.1 DUF4064 domain-containing protein
21(1) WP_185432669.1 DUF4064 domain-containing protein
21(1) WP_185387007.1 DUF4064 domain-containing protein
21(1) WP_196318787.1 DUF4064 domain-containing protein
21(1) WP_214169383.1 DUF4064 domain-containing protein
21(1) WP_185436221.1 DUF4064 domain-containing protein
21(1) WP_185639007.1 DUF4064 domain-containing protein
21(1) WP_185314064.1 hypothetical protein

22(2) WP_185365688.1 DUF1672 family protein
22(1) WP_185639010.1 DUF1672 family protein
22(1) WP_185489261.1 DUF1672 family protein
22(2) WP_243285278.1 hypothetical protein
22(1) WP_203577658.1 DUF1672 family protein
22(2) WP_185365689.1 DUF1672 family protein
22(1) WP_194360115.1 DUF1672 family protein
22(1) WP_185451612.1 DUF1672 family protein
22(1) WP_185639006.1 DUF1672 family protein

23(2) WP_228469892.1 glycohydrolase toxin TNT-related protein
23(2) WP_228469894.1 glycohydrolase toxin TNT-related protein
23(1) WP_203572938.1 glycohydrolase toxin TNT-related protein, partial
23(1) WP_185383808.1 glycohydrolase toxin TNT-related protein
23(1) WP_214169400.1 glycohydrolase toxin TNT-related protein
23(2) WP_185489269.1 hypothetical protein
23(1) WP_185469291.1 glycohydrolase toxin TNT-related protein
23(1) WP_185469293.1 pre-toxin TG domain-containing protein
23(1) WP_260443317.1 TNT domain-containing protein, partial

24(1) WP_185639020.1 WXG100 family type VII secretion target
24(1) WP_003728976.1 WXG100 family type VII secretion target
24(4) WP_011700921.1 WXG100 family type VII secretion target
24(3) WP_003745040.1 WXG100 family type VII secretion target
24(2) WP_012984597.1 WXG100 family type VII secretion target

25(1) WP_185639021.1 adenylosuccinate synthase
25(2) WP_011700920.1 adenylosuccinate synthase
25(1) WP_194361307.1 adenylosuccinate synthase
25(1) WP_185331645.1 adenylosuccinate synthase
25(1) WP_003728977.1 adenylosuccinate synthase
25(5) WP_003745033.1 adenylosuccinate synthase

26(5) WP_012984619.1 hypothetical protein
26(3) WP_185335964.1 hypothetical protein

27(8) WP_185336754.1 hypothetical protein

28(8) WP_185336755.1 hypothetical protein

29(3) WP_185365739.1 hypothetical protein
29(1) WP_194341478.1 hypothetical protein
29(2) WP_185383878.1 hypothetical protein
29(2) WP_185497587.1 hypothetical protein

30(1) WP_194341479.1 hypothetical protein
30(3) WP_185365741.1 hypothetical protein
30(3) WP_185497588.1 hypothetical protein
30(1) WP_185383810.1 hypothetical protein

31(2) WP_185383811.1 hypothetical protein
31(1) WP_194341480.1 hypothetical protein
31(3) WP_185365743.1 hypothetical protein
31(2) WP_185497590.1 hypothetical protein

32(4) WP_185383812.1 toxin B
32(3) WP_185365745.1 toxin B
32(1) WP_194341481.1 toxin B

33(1) WP_194321766.1 hypothetical protein
33(2) WP_185497593.1 hypothetical protein
33(1) WP_194341485.1 hypothetical protein
33(2) WP_185365749.1 hypothetical protein
33(2) WP_185436214.1 hypothetical protein

34(5) WP_185336756.1 Imm59 family immunity protein
34(2) WP_185326090.1 Imm59 family immunity protein

35(5) WP_012984617.1 DUF5085 family protein
35(1) WP_185455572.1 DUF5085 family protein
35(1) WP_203575362.1 DUF5085 family protein

36(1) WP_185606003.1 hypothetical protein
36(1) WP_185337277.1 hypothetical protein
36(2) WP_185415322.1 hypothetical protein
36(1) WP_228469158.1 hypothetical protein
36(2) WP_185432670.1 hypothetical protein

37(1) WP_194341477.1 DUF4176 domain-containing protein
37(2) WP_185365737.1 DUF4176 domain-containing protein
37(1) WP_194321764.1 DUF4176 domain-containing protein
37(2) WP_185497585.1 DUF4176 domain-containing protein

38(1) WP_185500481.1 hypothetical protein
38(2) WP_196318783.1 hypothetical protein
38(1) WP_185365700.1 hypothetical protein
38(1) WP_185387012.1 hypothetical protein
38(1) WP_185365698.1 hypothetical protein

39(4) WP_012984613.1 hypothetical protein

40(3) WP_012984612.1 DUF5082 family protein
40(1) WP_185489268.1 DUF5082 family protein

41(2) WP_012984614.1 T7SS effector LXG polymorphic toxin
41(1) WP_196318779.1 T7SS effector LXG polymorphic toxin
41(1) WP_185489267.1 T7SS effector LXG polymorphic toxin

42(1) WP_185365828.1 site-specific integrase
42(1) WP_228463714.1 tyrosine-type recombinase/integrase
42(1) WP_185455581.1 site-specific integrase

43(1) WP_228474721.1 hypothetical protein
43(1) WP_203577694.1 hypothetical protein
43(1) WP_185489285.1 hypothetical protein

44(2) WP_185313123.1 hypothetical protein

45(2) WP_185326091.1 helix-turn-helix transcriptional regulator

46(2) WP_185327600.1 hypothetical protein

47(2) WP_185327601.1 replication initiation factor domain-containing protein

48(2) WP_185415318.1 DUF5082 family protein

49(1) WP_185434086.1 hypothetical protein
49(1) WP_185415319.1 hypothetical protein

50(1) WP_185463557.1 CPBP family intramembrane metalloprotease
50(1) WP_185455583.1 CPBP family intramembrane metalloprotease

51(2) WP_228469893.1 IS110 family transposase

52(2) WP_228469905.1 transposase

53(1) WP_260445213.1 hypothetical protein
53(1) WP_185365826.1 hypothetical protein

Fig S4. Text output.
